# New insights into the functions of ACBD4/5-like proteins using a combined phylogenetic and experimental approach across model organisms

**DOI:** 10.1101/2024.06.21.599987

**Authors:** Suzan Kors, Martin Schuster, Daniel C. Maddison, Sreedhar Kilaru, Tina A. Schrader, Joseph L. Costello, Markus Islinger, Gaynor A. Smith, Michael Schrader

## Abstract

Acyl-CoA binding domain-containing proteins (ACBDs) perform diverse but often uncharacterised functions linked to cellular lipid metabolism. Human ACBD4 and ACBD5 are closely related peroxisomal membrane proteins, involved in tethering of peroxisomes to the ER and capturing fatty acids for peroxisomal β-oxidation. ACBD5 deficiency causes neurological abnormalities including ataxia and white matter disease. Peroxisome-ER contacts depend on an ACBD4/5-FFAT motif, which interacts with ER-resident VAP proteins. As ACBD4/5-like proteins are present in most fungi and all animals, we combined phylogenetic analyses with experimental approaches to improve understanding of their evolution and functions. Notably, all vertebrates exhibit gene sequences for both ACBD4 and ACBD5, while invertebrates and fungi possess only a single ACBD4/5-like protein. Our analyses revealed alterations in domain structure and FFAT sequences, which help understanding functional diversification of ACBD4/5-like proteins. We show that the *Drosophila melanogaster* ACBD4/5-like protein possesses a functional FFAT motif to tether peroxisomes to the ER via Dm_Vap33. Depletion of Dm_Acbd4/5 caused peroxisome redistribution in wing neurons and reduced life expectancy. In contrast, the ACBD4/5-like protein of the filamentous fungus *Ustilago maydis* lacks a FFAT motif and does not interact with Um_Vap33. Loss of Um_Acbd4/5 resulted in an accumulation of peroxisomes and early endosomes at the hyphal tip. Moreover, lipid droplet numbers increased and mitochondrial membrane potential declined, implying altered lipid homeostasis. Our findings reveal differences between tethering and metabolic functions of ACBD4/5-like proteins across evolution, improving our understanding of ACBD4/5 function in health and disease. The need for a unifying nomenclature for ACBD proteins is discussed.

## 1. Introduction

Acyl-CoA binding domain-containing proteins (ACBDs) comprise a large multigene family of diverse proteins with a conserved acyl-CoA binding motif, and are found in animals, plants and fungi, as well as in bacteria and archaea (Neess et al., 2015; Islinger et al., 2020). Activated fatty acids (acyl-CoAs) play important roles as lipid metabolites, but also in the regulation of lipid metabolism and in cellular signalling, and ACBD proteins fulfil important functions in controlling their concentration (Færgeman and Knudsen, 1997; Islinger et al., 2020). Notably, ACBDs are now also recognised as targets for different pathogens including viruses, *Salmonella* and *Chlamydia* (Islinger et al., 2020). Despite their fundamental importance to cellular lipid metabolism, the functions of many ACBD proteins remain unclear. In mammals, eight ACBD proteins (ACBD1-8) have been identified, with several (ACBD2, ACBD4, ACBD5) having been linked to peroxisome function (Costello et al., 2017b; c, 2023; Fan et al., 2016). Peroxisomes are oxidative organelles with essential functions in cellular lipid metabolism, including fatty acid α- and β-oxidation and the synthesis of ether-phospholipids (e.g. in myelin sheaths). Many of the peroxisomal lipid-metabolising functions are performed in cooperation with other subcellular organelles such as mitochondria (e.g. fatty acid β-oxidation) and the ER (e.g. ether lipid biosynthesis) (Wanders et al., 2018).

ACBD4 and ACBD5 are C-terminally tail-anchored peroxisomal membrane proteins, which expose their N-terminal acyl-CoA binding domain to the cytosol (Costello et al., 2017a). We recently revealed that ACBD4 and ACBD5 can act as peroxisome-ER tethers through binding of ER-resident VAPs (vesicle-associated membrane protein [VAMP]-associated proteins A/B). The interaction is mediated by a FFAT motif (two phenylalanines [FF] in an acidic tract) in the central region of ACBD4 and ACBD5, which binds to the MSP (major sperm protein) domain of VAPs (Costello et al., 2017b; c, 2023; Hua et al., 2017) (**Fig. 1A**). This interaction is regulated by phosphorylation of the ACBD5 FFAT motif, but similar regulation of ACBD4 was not observed (Kors et al., 2022b). Whilst both proteins have the capacity to act as tethers, our recent work suggested that ACBD5 may play a more significant part than ACBD4 in binding VAPB to provide a physical tether, whereas ACBD4 may localise to the peroxisome-ER contact sites to provide a regulatory function (Costello et al., 2023). ACBD5-VAP interactions create peroxisome-ER membrane contact sites that are important for lipid transfer to support cooperative metabolism and peroxisome biogenesis (Schrader et al., 2020; Carmichael et al., 2022).

**Fig. 1.**
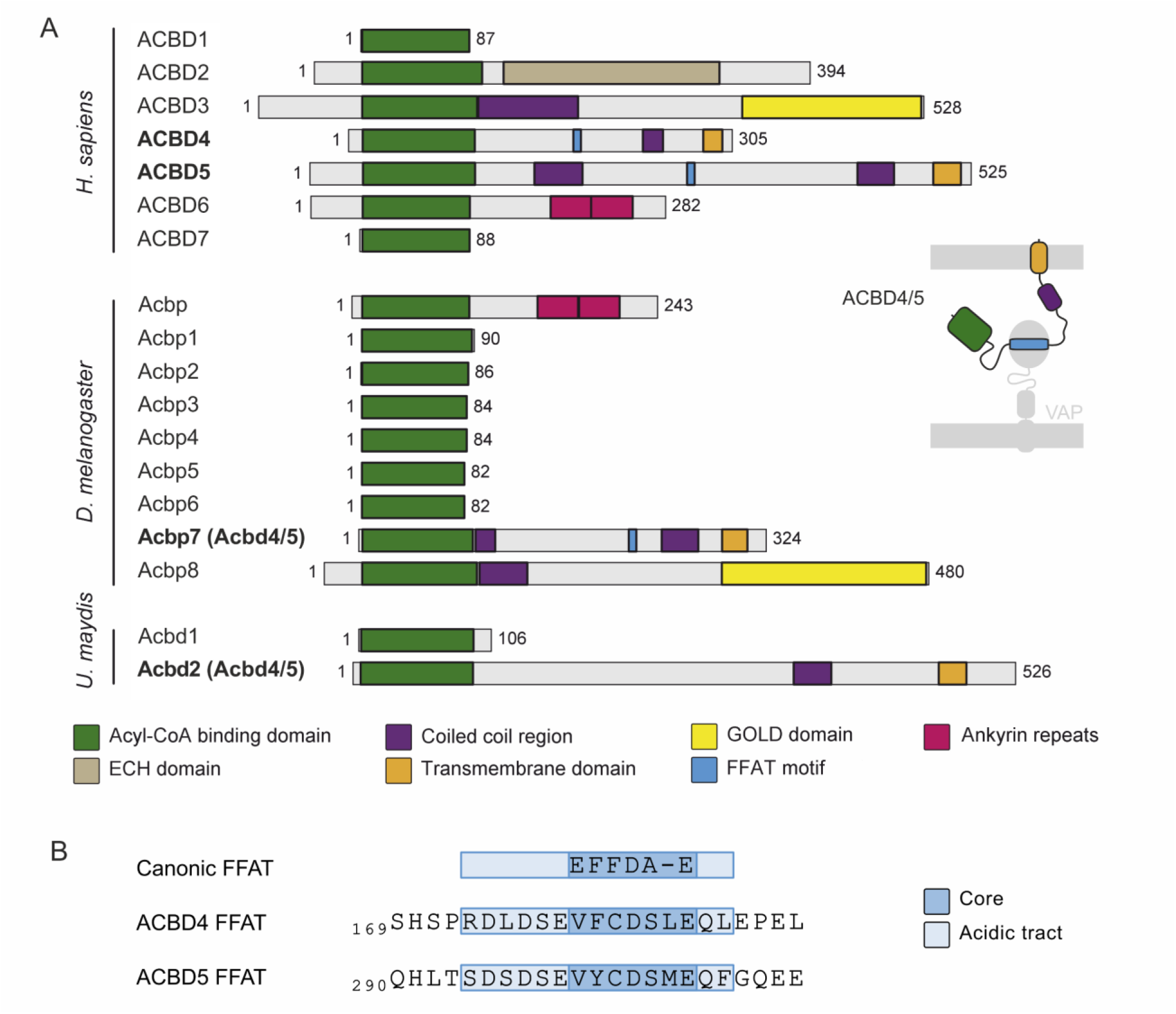
Domain structures of human, fly and fungal ACBD proteins. **(A)** The schemes represent ACBDs from *Homo sapiens* (Mammalia, Animalia), *Drosophila melanogaster* (Insecta, Animalia) and *Ustilago maydis* (Basidiomycota, Fungi). Sequence and domain lengths as well as position are proportional and aligned to the acyl-CoA binding (ACB) domains. In *H. sapiens* seven ACBD proteins have been identified: ACBD1 (ACBP, DBI), ACBD2 (ECI2, PECI), ACBD3 (GCP60, PAP7), ACBD4, ACBD5, ACBD6 and ACBD7. The *D. melanogaster* genome has a total of nine genes encoding putative ACBDs: Acbp (Anox, Acbp-Ank, CG33713), Acbp1 (CG8498), Acbp2 (Dbi, CG8627), Acbp3 (CG8628), Acbp4 (CG8629), Acbp5 (CG5804), Acbp6 (CG15829), Acbp7 (Acbd4/5, CG8814), and Acbp8 (CG14232); and *U. maydis* two: Acbd1 (UMAG_02959) and Acbd2 (Acbd4/5, UMAG_11226). For some ACBDs, several transcript variants have been reported or predicted. ECH, enoyl-CoA hydratase; FFAT, two phenylalanines in an acidic tract motif; GOLD, Golgi dynamics. **(B)** Amino acid sequences of the FFAT motif of human ACBD4 (Q8NC06-2) and ACBD5 (Q5T8D3-3).

In addition to their tethering role, ACBD4 and ACBD5 may have a metabolic function in capturing fatty acids with their acyl-CoA binding motif for peroxisomal β-oxidation (Costello et al., 2023). ACBD5 has a preference for very long chain-fatty acids (VLCFA) and is supposed to capture them for hand-over to the peroxisomal VLCFA transporter ABCD1, which imports them for subsequent peroxisomal fatty acid β-oxidation. Peroxisome tethering to the ER is thought to create a lipid hub, which allows tight coordination of fatty acid synthesis and elongation at the ER and breakdown by peroxisomal β-oxidation according to metabolic needs (Schrader et al., 2020).

Meanwhile, numerous patients with an ACBD5 deficiency have been identified which present with retinal dystrophy, ataxia, psychomotor delay and a severe leukodystrophy (Carmichael et al., 2022; Abu-Safieh et al., 2013; Ferdinandusse et al., 2017; Yagita et al., 2017; Helman et al., 2020; Bartlett et al., 2021; Gorukmez et al., 2022; Pappaterra-Rodriguez et al., 2022; Hasturk et al., 2024; Rudaks et al., 2024). Loss of ACBD5 leads to an accumulation of VLCFA, due to impaired VLCFA import/β-oxidation in peroxisomes, as well as a reduction in ether-phospholipids (Herzog et al., 2018), which are synthesised cooperatively by peroxisomes and the ER. Two *Acbd5* knockout mouse models have been generated which, like human patients, show an increase in VLCFA and develop a neurological pathology characterised by a progressive motor dysfunction and retinal degeneration (Darwisch et al., 2020; Granadeiro et al., 2023).

Although structurally and phylogenetically closely related, ACBD4 and ACBD5 display differences in substrate binding, regulation, tethering capacity and expression (Costello et al., 2023). ACBD4 appears to have evolved from ACBD5 by gene duplication, which only occurred in vertebrates (Islinger et al., 2020). Our recent phylogenetic analysis of the ACBD protein family revealed that homologs of ACBD4/5 are present in many fungi and metazoans, but absent in, for instance, *Saccharomyces cerevisiae* (Camões et al., 2015; Islinger et al., 2020). Furthermore, according to bioinformatics analysis, not all ACBD4/5-like proteins contain a FFAT motif (Islinger et al., 2020), and therefore the contribution of ACBD4 and ACBD5 to peroxisome-ER tethering might not be a general and original feature/function of the proteins.

To understand the evolution and function of ACBD4/5 proteins in more detail, we have investigated them in two easily tractable model organisms, the fruit fly *Drosophila melanogaster* and the filamentous fungus *Ustilago maydis. D. melanogaster* has been an important model organism for many human diseases, including neurological disorders (Pandey and Nichols, 2011). For instance, adult wing neurons are easily accessible for studying axonal organelle distribution (Smith et al., 2019). Furthermore, many human peroxisomal proteins and pathways are conserved in the fly (Faust et al., 2012; Baron et al., 2016). Several studies have shown that mutations in peroxisomal biogenesis genes cause similar phenotypes as seen in humans (Mast et al., 2011), making *D. melanogaster* a useful model to study peroxisomal disorders.

*U. maydis* is a phytopathogenic fungus, causing smut disease in maize (*Zea mays*). The basidiomycetous fungus has been introduced as a new model organism to study basic cell biological processes (Steinberg and Perez-Martin, 2008). Its filamentous appearance (hyphal cells) and technical advantages provide a system to study long distance intracellular transport, polarised growth and motor-based microtubule organisation – key processes in neuronal cells. Furthermore, many peroxisomal proteins and pathways are shared between *U. maydis* and humans (Camões et al., 2015). Whereas in yeast and plants, fatty acids are only oxidised in peroxisomes, *U. maydis* and humans have cooperative peroxisomal and mitochondrial β-oxidation pathways.

Here, we combined phylogenetic analysis of ACBD4/5-like proteins in animals and fungi with experimental approaches. We revealed that the Acbd4/5 protein in *D. melanogaster* possesses a functional FFAT motif to tether peroxisomes to the ER via Vap33 binding. Depletion of Acbd4/5 in *Drosophila* leads to peroxisome redistribution in wing neurons and reduced life expectancy. In contrast, *U. maydis* Acbd4/5 does not possess a FFAT motif and does not interact with Vap33. Although loss of the fungal Acbd4/5 does not impact on cellular growth, it results in impaired distribution of peroxisomes and early endosomes in hyphae. Differences between the tethering and metabolic functions of ACBD4/5-like proteins are discussed.

## 2. Results

### 2.1 Domain comparison of the ACBD4/5 homologs in H. sapiens, U. maydis and D. melanogaster

Humans (Hs) encode seven ACBD proteins (ACBD1-7), with diverse subcellular location and functions (Islinger et al., 2020). Our recent phylogenetic analysis of the ACBD protein family, defined by the presence of an acyl-CoA binding motif (ACB), showed that the ACBD proteins are highly conserved across species (Islinger et al., 2020). BLAST searches revealed that *D. melanogaster* and *U. maydis* encode nine and two putative ACBD proteins, respectively (**Fig. 1A**), with one in each species showing similar domain organisation to the closely related peroxisomal membrane proteins Hs_ACBD4 and Hs_ACBD5.

An expanded, more focused phylogenetic analysis of 275 ACBD4/5-like sequences from 189 animal and fungal species revealed that all animals and most fungi contain ACBD4/5-like sequences with a highly conserved ACB domain, a coiled-coil region in the intermediate part and a transmembrane domain (TMD) with an adjacent stretch of positively charged amino acids at the C-terminus (**Fig. 1A**). Notably, all vertebrates, including the agnathous lamprey *Petromyzon marinus*, exhibit sequences for both ACBD4 and ACBD5, while invertebrates and fungi possess only a single ACBD4/5-like protein.

Assessing the ACBD4/5 sequences in evolutionary distant vertebrates may assist to delineate how a single precursor differentiated into the two mammalian proteins. Human ACBD4 is a significantly more compact protein than human ACBD5, which lacks the extended sequence connecting the ACB domain with a coiled-coiled region adjacent to the C-terminal TMD (**Fig. 1A**). This generally largely distorted region contains a second predicted coiled-coil motif, enriched in negatively charged amino acids. Of note, fish ACBD4 sequences get, in evolutionary terms, continuously longer in more “primitive” species like *Chondrichthyes* or *Agnatha* showing homologies across the whole protein sequence.

Mammalian ACBD4 and ACBD5 both contain a FFAT motif, which consists of seven core residues flanked by an acidic tract, with which they bind to ER-resident VAP proteins to mediate peroxisome-ER contacts (Costello et al., 2017b; c) (**Fig. 1B**). However, ACBD5 does this more efficiently than ACBD4 (Kors et al., 2022b; Costello et al., 2023), which is also documented by the significantly stronger predicted FFAT scores found in mammalian ACBD5 compared to ACBD4 sequences (x vs y; **Table 1, Fig. S1, Sup. Data 1**), as determined by a FFAT motif prediction tool (Murphy and Levine, 2016). Mutations in the N-terminal region of the ACBD4 FFAT motif appear to have modified the VAP-binding properties of the protein. While the ACBD5 FFAT motif is characterised by a conserved negatively charged SDSDS acidic tract sequence in front of the hydrophobic V/Y/F core residues, ACBD4-sequences from mammals and birds exhibit amino acid substitutions, which alter the polarity of the acidic tract (**Fig. 2A**). Thus, after an ACBD4/5 gene duplication occurred at the base of the vertebrate branch, ACBD4 appears to have evolved from ACBD4/5 by consecutively reducing its tethering function. In parallel, in the ACB domain a substitution of a conserved glutamine (Q) by an arginine (R) in the region between helix 1 and helix 2 as well as an insertion of four amino acids at the C-terminus of helix 4 may have changed the substrate binding properties (**Fig. 2B**).

**Fig. 2.**
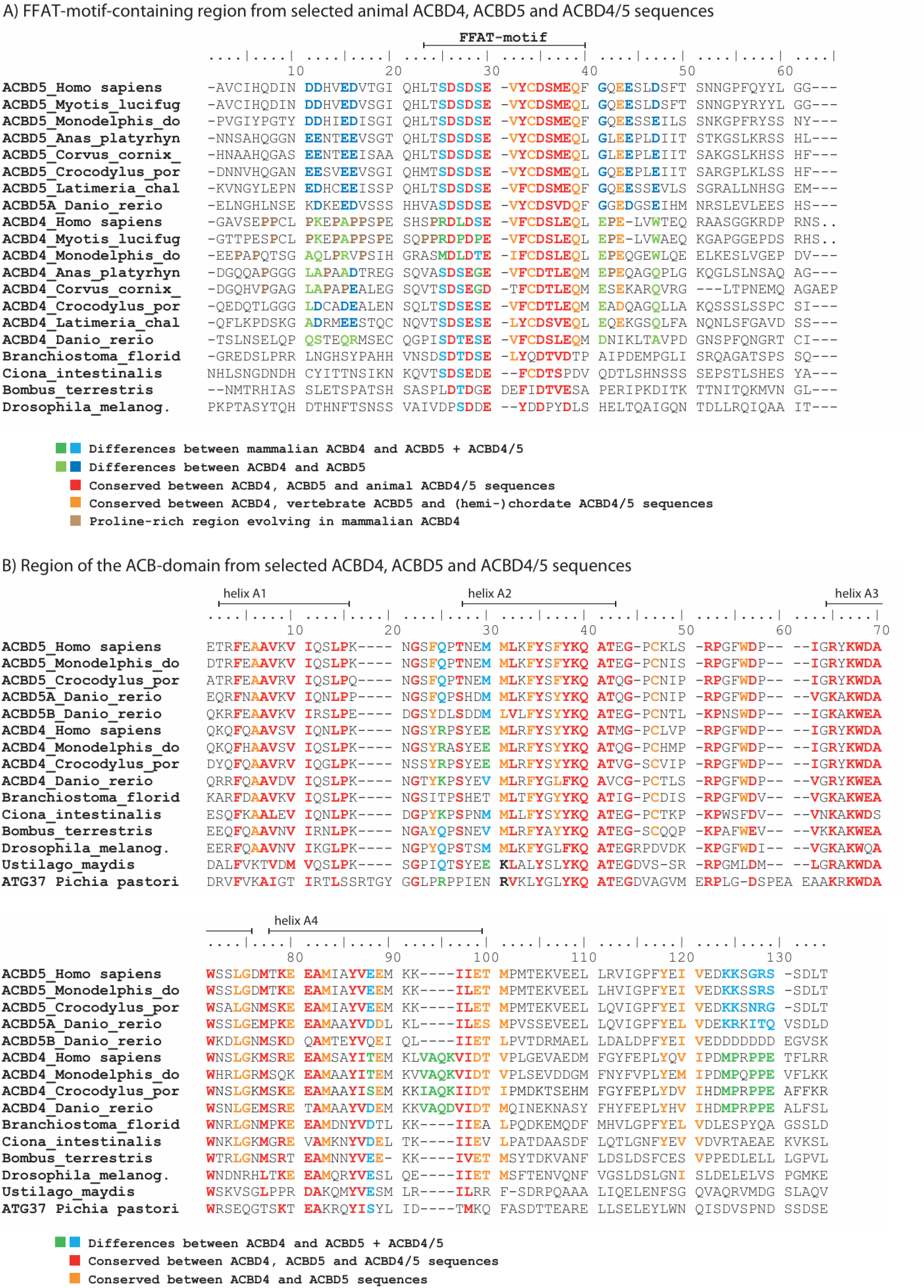
Conservation of the ACB domain and the FFAT motif in ACBD4/5 sequences. **(A)** Amino acid conservation in the region of the FFAT motif from selected animal ACBD4 and ACBD5 sequences. Amino acids with equal physicochemical properties conserved among all animal groups are highlighted in **red**, those conserved only in vertebrate ACBD5 and ACBD4 sequences in **orange**. Amino acids, which show significant sequence differences between ACBD4 and ACBD5 sequences are highlighted in **green** vs. **blue**. A proline-rich region N-terminal to the FFAT motif, which is specifically found in mammalian species, is highlighted in **brown**. **(B)** Alignment of the ACB domains of representative vertebrate, invertebrate and fungal ACBD4/5 sequences. Amino acids with equal physicochemical properties conserved among all phylogenetic groups are highlighted in **red**, those conserved only in vertebrate ACBD5 and ACBD4 sequences in **orange**. Amino acids, which show significant sequence variation among subgroups or species clusters, are highlighted in **green** vs. **blue**.

**Table 1.** Potential FFAT motifs in metazoan ACBD4/5-like sequences.

|  | Ø FFAT motif score | potential phosphorylation | conserved position |
| --- | --- | --- | --- |
| ACBD4/5<br>Basidiomycota | 4.7 | no | no |
| ACBD4/5<br>Ascomycota | 4.3 | no | no |
| ACBD4/5<br>Other fungi | 4.1 | no | no |
| ACBD5<br>Mammalia | 2.5 | yes | yes |
| ACBD5<br>Aves | 2.5 | yes | yes |
| ACBD5<br>Reptilia | 2.5 | yes | yes |
| ACBD5<br>Amphibia | 2.5 | yes | yes |
| ACBD5<br>Osteichthyes | 2.5 | yes | yes |
| ACBD5<br>Chondrichthyes | 2.4 | yes | Yes |
| ACBD5<br>Agnatha | 2.5 | yes | yes |
| ACBD5B<br>Teleostei | 2.8 | partially | partially |
| ACBD4<br>Mammalia | 3.4 | yes<br>except *Tachyglossus* | yes<br>except *Tachyglossus* |
| ACBD4<br>Aves | 2.4 | yes | yes |
| ACBD4<br>Reptilia | 2.6 | yes | yes |
| ACBD4<br>Amphibia | 2.3 | partially | yes |
| ACBD4<br>Osteichthyes | 2.6 | yes<br>except *Polypterus* | yes<br>except *Polypterus* |
| ACBD4<br>Chondrichthyes | 2.5 | yes | yes |
| ACBD4<br>Agnatha | 2.5 | yes | yes |

ACBD4/5 FFAT motifs can be strengthened or weakened by phosphorylation of conserved serine/threonine residues inside the acidic tract or the FFAT core, respectively (Kors et al. 2022b). “Potential phosphorylation” indicates the conservation of these residues in the FFAT motif.
“Conserved position” indicates that the FFAT motif is positioned at a conserved site inside the proteins primary sequence.

As shown in a phylogenetic cladogram, *D. melanogaster* CG8814 (hereafter Dm_Acbd4/5) and *U. maydis* UMAG_11226 (hereafter Um_Acbd4/5) may be orthologs for human ACBD4 and ACBD5 (**Fig. S1; Fig. 1A**). We set out to decipher if the insect and basidiomyocote proteins share more features with either ACBD4 or ACBD5, combine sequence motifs and functions from both mammalian proteins, or have developed their own unique characteristics. Compared to the vertebrate ACBD5, insect Acbd4/5 sequences including *D. melanogaster* Acbd4/5 are in average significantly shorter and, hence, with respect to their general structure, closer to mammalian ACBD4. In general, arthropod Acbd4/5 sequences possess a typical FFAT motif with a strong acidic tract (e.g. *Bombus terrestris* in **Fig. 2A**). In *Drosophila* species, however, the serine/threonine (S/T) in the core of the FFAT motif was replaced by a proline (P), which could considerably weaken the FFAT motif (**Fig. 2A**).

In contrast to the short ACBD4/5 sequences from insects, fungal Acbd4/5 sequences vary substantially in size. While the ACBD4/5-like protein Atg37 in *Pichia pastoris* (Nazarko et al., 2014) has a sequence length of 409 amino acids, which is closer to Hs_ACBD4, *U. maydis* Acbd4/5 contains 526 amino acids, which is more similar to Hs_ACBD5. A coiled-coil region, involved in the dimerisation of human ACBD5 (Costello et al., 2023), is also present in Um_Acbd4/5. However, compared to the human proteins, the Um_Acbd4/5 coiled-coil region, which is generally present in the fungal proteins, lies further towards the C-terminus resulting in a longer ’gap’ upstream of the TMD (**Fig. 1A**). Additionally, fungal Acbd4/5 sequences including Um_Acbd4/5 exhibit a longer tail-sequence downstream of the TMD, which therefore localises inside peroxisomes and exhibits an additional conserved short α-helical segment.

Both fungal as well as all animal ACBD4/5 sequences are conserved in all residues of the ACB domain described to be required for the binding of long-chain acyl-CoAs (Kragelund et al., 1999; van Aalten et al., 2001). As expected, a higher degree of sequence variation becomes obvious when *U. maydis* Acbd4/5 or *P. pastoris* Atg37 are compared with animal ACBD4 and ACBD5 sequences, which might result in slight differences in the binding spectrum of acyl-CoAs. Interestingly, a highly conserved methionine (M) from helix 2 in animals was replaced by lysine (K) or arginine (R) in fungi (**Fig. 2B**). Of note, the same sequence difference was reported between mammalian and *Plasmodium* ACBP (ACBD1) (van Aalten et al., 2001). Since the exchange adds an additional positive charge to the interior of the acyl-CoA binding pocket, the authors suggested that additional acceptor molecules with negative charges might be used to block the binding pocket, adding another possibility for regulating substrate binding. Most importantly, we were not able to identify any potential FFAT motif in Um_Acbd4/5. This is in line with our analysis of the presence of FFAT motifs in ACBD4/5-like proteins across species, where we identified FFAT motifs in almost all animal ACBD4/5 homologs (95%), but not in the majority of the fungal homologs analysed (20%).

### 2.2 Peroxisomal targeting of D. melanogaster and U. maydis Acbd4/5

Targeting of the human ACBD4 and ACBD5 proteins to peroxisomes depends on the hydrophobicity and charge of their transmembrane and tail region (Costello et al., 2017a; b; c; Hua et al., 2017). Using our previously developed targeting algorithm for C-tail-anchored membrane proteins (Costello et al., 2017a), we predicted that *D. melanogaster* and *U. maydis* Acbd4/5 would localise to peroxisomes. To confirm peroxisomal targeting, Dm_Acbd4/5 and Um_Acbd4/5 were cloned into a mammalian expression vector and expressed in COS-7 cells. Both Dm_Acbd4/5 and Um_Acbd4/5 colocalised with the peroxisomal membrane marker PEX14 (**Fig. 3A, B**), indicating conserved physicochemical properties for the peroxisomal targeting of the ACBD4/5 homologs. Indeed, peroxisomal targeting of Acbd4/5 was recently confirmed in *D. melanogaster* (Barone et al., 2023a). Interestingly, expression of Dm_Acbd4/5, but not Um_Acbd4/5, induced peroxisome elongation in COS-7 cells (**Fig. 3A, B**), similar to overexpression of Hs_ACBD5 (Costello et al., 2017b; Kors et al., 2022b). Peroxisome elongation and membrane expansion have been linked to the peroxisome-ER tethering function of ACBD5, which is required for membrane lipid transfer. Overall, these findings suggest that the Hs_ACBD5 and Dm_Acbd4/5 proteins are functional orthologs, whereas Um_Acbd4/5 and Hs_ACBD5 appear to be sequence homologs, which may have acquired different peroxisome-associated functions.

**Fig. 3.**
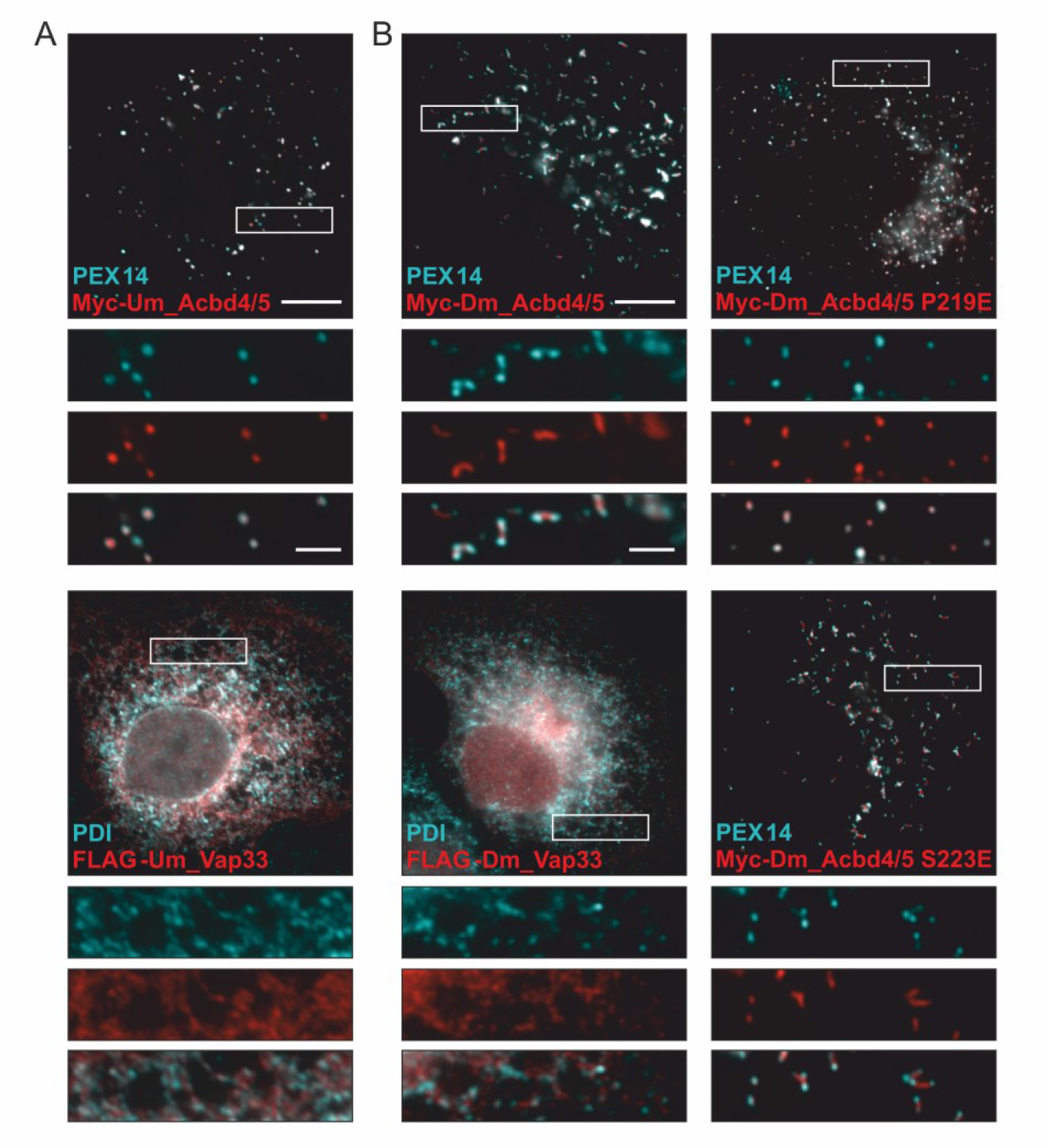
Subcellular localization of the *U. maydis* and *D. melanogaster* ACBD4/5 and VAP homologs. COS-7 cells transfected with **(A)** *U. maydis* (Um) Myc-Acbd4/5 or FLAG-Vap33, or **(B)** *D. melanogaster* (Dm) Myc-Acbd4/5 (WT, S219E and S223E) or FLAG-Vap33, were immunolabeled with PEX14 (peroxisomal membrane marker), PDI (ER lumen marker) and Myc/FLAG antibodies. Bars: 10 µm (main), 2.5 µm (inset).

### 2.3 D. melanogaster Acbd4/5 binds to Vap33 via a FFAT motif

*D. melanogaster* has one highly conserved VAP homolog, Vap33, which shares the domain organisation of human VAPA/B (Chai et al., 2008) (**Fig. 4A**). This suggests that Acbd4/5 could bind Vap33 via its predicted FFAT motif and mediate peroxisome-ER contacts, as in humans. To test this, Dm_Vap33 was cloned into a mammalian expression vector and expressed in COS-7 cells, where it showed ER-targeting (**Fig. 3B**). Acbd4/5-Vap33 binding was confirmed by co-immunoprecipitation after co-expression of Myc-Dm_Acbd4/5 and FLAG-Dm_Vap33 in COS-7 cells (**Fig. 4B**). Supporting this, Acbd4/5 was identified as a potential binding partner of Vap33 in a study that performed high-throughput mapping of the fly interactome using affinity purification-mass spectrometry (Guruharsha et al., 2011).

**Fig. 4.**
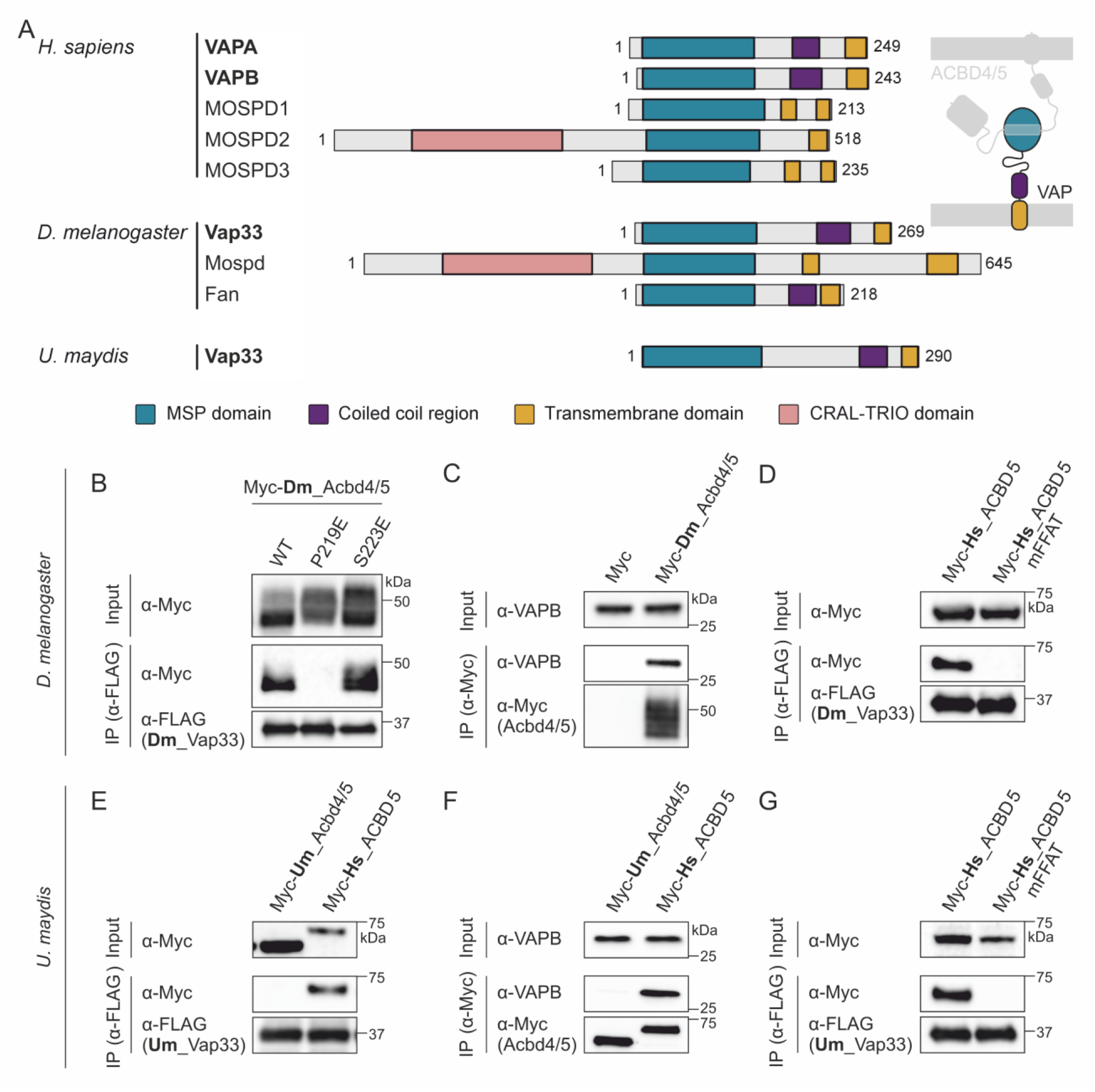
*D. melanogaster* Acbd4/5, but not *U. maydis* Acbd4/5, binds to Vap33 via a FFAT motif. **(A)** The schemes represent VAPs from *Homo sapiens* (Mammalia, Animalia), *Drosophila melanogaster* (Insecta, Animalia) and *Ustilago maydis* (Basidiomycota, Fungi). Sequence and domain lengths as well as position are proportional and aligned to the major sperm protein (MSP) domains. In *H. sapiens* five VAP proteins have been identified: VAPA (VAP33), VAPB, MOSPD1, MOSPD2 and MOSPD3. *D. melanogaster* has three VAPs: Vap33 (dVap-A, CG5014), Mospd (dVap-B, CG33523) and Fan (farinelli, dVap-C, CG7919 – testis specific); and *U. maydis* has one putative VAP: Vap33 (UMAG_11696). For some VAPs, several isoforms have been reported or predicted. CRAL-TRIO, cellular retinaldehyde-binding protein and triple functional domain protein; MOSPD, motile sperm domain-containing protein; VAP, vesicle-associated membrane protein (VAMP)-associated protein. **(B-G)** Binding assays with *D. melanogaster* (Dm), *U. maydis* (Um) and *H. sapiens* (Hs) constructs expressed in COS-7 cells. Samples were used to perform immunoprecipitation (IP), and subsequently immunoblotted using Myc/FLAG/VAPB antibodies. **(B)** IP with FLAG-Dm_Vap33 to detect binding of Myc-Dm_Acbd4/5. Myc-Dm_Acbd4/5 P219E and S223E constructs contain a mutation of a critical residue in the predicted FFAT motifs. **(C)** IP with Myc-Dm_Acbd4/5 (or control vector [Myc]) to detect bound endogenous VAPB. **(D)** IP with FLAG-Dm_Vap33 to detect binding of Myc-Hs_ACBD5 WT or with a mutated FFAT motif (mFFAT). **(E)** IP with FLAG-Um_Vap33 to detect binding of Myc-Um_Acbd4/5. The confirmed VAP interactor Myc-Hs_ACBD5 was used as positive control. **(F)** IP with Myc-Um_Acbd4/5 (or Myc-Hs_ACBD5 as positive control) to detect bound endogenous VAPB. **(G)** IP with FLAG-Um_Vap33 to detect binding of Myc-Hs_ACBD5 WT or mFFAT.

To determine whether Acbd4/5 binds Vap33 via a FFAT motif, we generated Myc-Dm_Acbd4/5 in which a critical residue in the predicted FFAT motifs was mutated (P219E) (Kors et al., 2022b). Interestingly, *D. melanogaster* Acbd4/5 exhibits, in addition to the predicted _2151_EYDDPYD^7^ FFAT motif, a potential second, overlapping FFAT _2191_PYDLSHE^7^, which we tested in parallel (S223E). Both mutants were properly targeted to peroxisomes (**Fig. 3B**). The Acbd4/5-Vap33 interaction was lost with the Dm_Acbd4/5 P219E mutant, whereas Dm_Acbd4/5 S223E was not affected in its binding to Dm_Vap33. The FFAT-MSP interaction was conserved across species: Dm_Acbd4/5 bound to endogenous mammalian VAPB (**Fig. 4C**), and Dm_Vap33 interacted with Hs_ACBD5, which was abolished when the FFAT motif was mutated (Hs_ACBD5 mFFAT) (**Fig. 4D**). In conclusion, *D. melanogaster* Acbd4/5 and Vap33 interact via the Acbd4/5 FFAT motif _2151_EYDDPYD^7^, which also corresponds to the position of the FFAT motif in the mammalian ACBD4 and ACBD5 proteins in the sequence alignment (**Fig. 2A**).

Although the fungal Um_Acbd4/5 lacks a predicted FFAT motif, an additional BLAST search revealed that *U. maydis* does contain a VAP homolog, UMAG_11696 (hereafter Vap33), with a conserved MSP domain (**Fig. 4A**). Um_Vap33 was cloned into a mammalian expression vector and showed ER targeting when expressed in COS-7 cells (**Fig. 3A**). However, as predicted, Myc-Um_Acbd4/5 was not immunoprecipitated by FLAG-Um_Vap33 (**Fig. 4E**) or FLAG-Hs_VAPB (**Fig. 4F**), indicating that *U. maydis* Acbd4/5 does not interact with Vap33 and has no FFAT motif that can mediate binding to Vap33/VAPB.

Next, we assessed if *U. maydis* Vap33 has the ability to bind FFAT motifs using human ACBD5. We observed that FLAG-Um_Vap33 immunoprecipitated with Myc-Hs_ACBD5, but not Myc-Hs_ACBD5 mFFAT (**Fig. 4G**). This implies that Um_Vap33 can nevertheless bind FFAT motifs with its MSP domain. Like human VAPA/B, Um_Vap33 contains a C-terminal TMD and is therefore theoretically able to act as a tethering protein at membrane contact sites. We screened homologs of other known human FFAT-containing proteins in *U. maydis* and discovered that unlike Acbd4/5, UMAG_11568 (ORP3-related), Osh3 and Vps13 contain predicted FFAT motifs, so may represent Vap33 interacting partners. Overall, these experiments show that *H. sapiens* and *D. melanogaster* ACBD5/Acbd4/5 interact with VAPB/Vap33 via their FFAT motif, while the *U. maydis* homologs Acbd4/5 and Vap33 do not interact.

### 2.4 Depletion of Acbd4/5 or Vap33 disrupts peroxisome distribution in D. melanogaster wing neurons

Silencing of ACBD5 or both VAPA and VAPB in mammalian cells leads to increased peroxisome motility (Costello et al., 2017b; Hua et al., 2017), and overexpression of ACBD5 was shown to reduce peroxisomal long-range movements in the neurites of mouse primary hippocampal neurons (Wang et al., 2018). Therefore, we hypothesised that disrupting peroxisome-ER interaction by ACBD5 or VAP depletion could result in more mobile peroxisomes translocating into neurites and changing peroxisome positioning. We explored this using Acbd4/5 and Vap33 in *D. melanogaster* as a model. In order to visualise peroxisomes in neurons, we used flies expressing GFP fused to a peroxisomal targeting signal (GFP-SKL) and tdTomato (5x*UAS-SKL::GFP, 10xUAS-IVS-myr::tdTomato*) under the control of a pan-neuronal driver (*nSyb-GAL4*) to label the peroxisomal matrix and cell membrane, respectively, in mature adult neurons. These flies were crossed with flies harbouring either a heterozygous mutant *acbd4/5* (*acbd4/5^+/-^*) or an *acbd4/5* RNAi transgene (*5xUAS-acbd4/5^RNAi^*) to examine the effect of *acbd4/5* depletion on peroxisome distribution in the easily accessible wing neurons. Neurons proximal and distal to the fly body were imaged at 1 and 7 days post-eclosion (DPE; i.e. the time from when adults emerge from their pupal case) (**Fig. 5A**). In general, peroxisomes were predominantly located in the cell body, with a low number of peroxisomes found in neurites (**Fig. 5B, C**). The extended morphology and positioning of the long sensory axon tracts allowed us to quantify the number of individual peroxisomes located in these projections compared to the somato-dendritic region. An increase in the number of axonal peroxisomes in *acbd4/5* depleted flies was observed compared to control, in both proximal and distal neurons (**Fig. 5D, E**). To exclude that this rise in axonal peroxisomes could be caused by an increase in peroxisome number, we quantified the total cellular peroxisome content in the analysed neurons. Importantly, no difference between the cellular peroxisome content of the *acbd4/5* depletion and control lines was detected (**Fig. 5F, G**). In general, the cellular peroxisome content decreased in ageing flies (**Fig. 5H, I**), but this did not seem to significantly affect axonal peroxisome number in the control flies between 1 and 7 DPE (**Fig. 5D, E**).

**Fig. 5.**
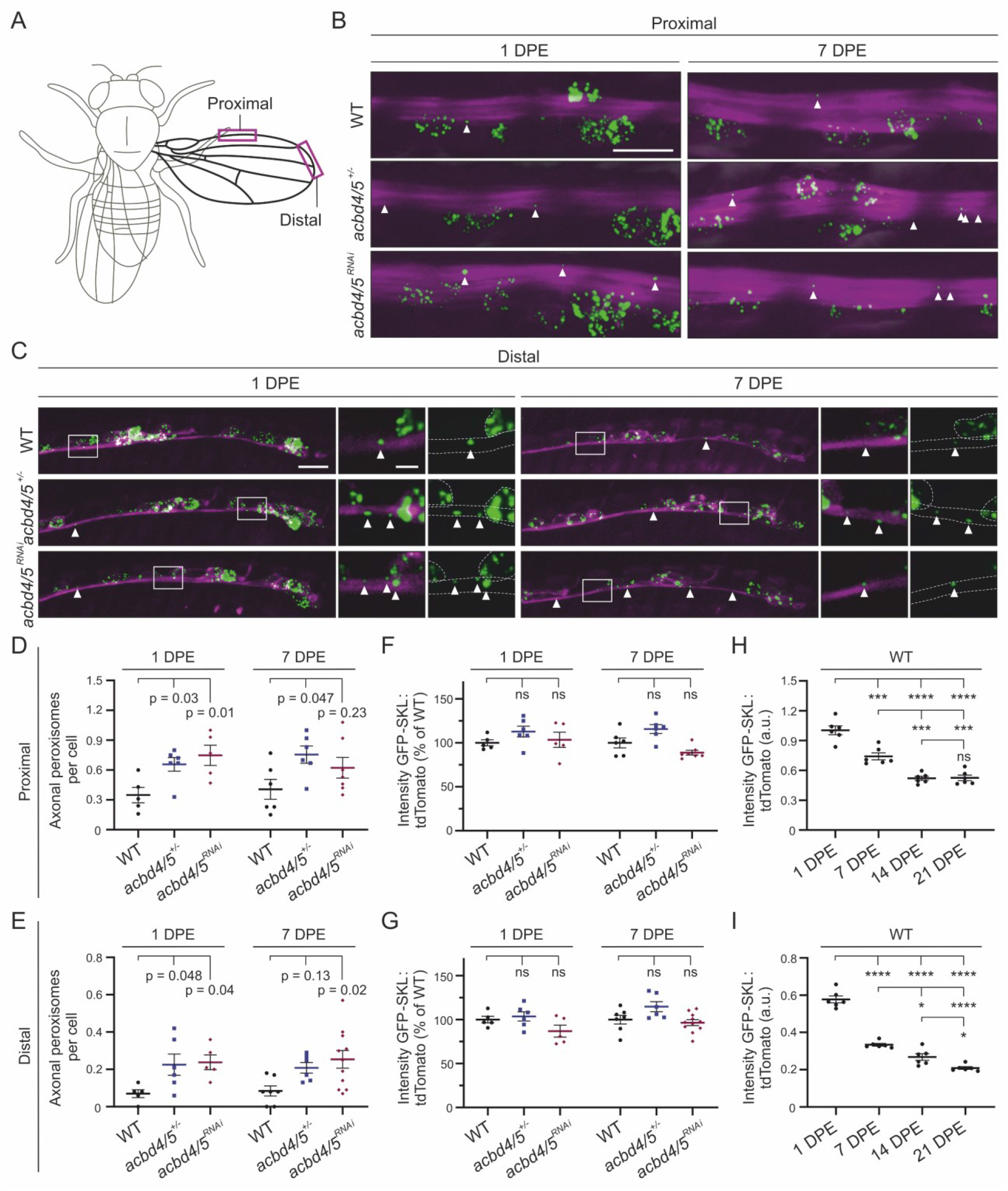
Depletion of *acbd4/5* in *D. melanogaster* increases the number of axonal peroxisomes. **(A)** The imaged areas to study peroxisomes in both proximal and distal wing neurons are mapped on the fly. **(B-C)** Flies expressing GFP-SKL and tdTomato to label peroxisomes and the cytosol, respectively, in mature neurons were used (*nSyb-GAL4, UAS-GFP::SKL, UAS-tdTomato*). Proximal **(B)** and distal **(C)** neurons were imaged at 1 and 7 days post-eclosion (DPE) in *acbd4/5* depleted (*acbd4/5^+/-^*and *acbd4/5^RNAi^*) flies. Insets present selected planes of the Z-projection shown in the main image. **(D-E)** The number of peroxisomes located in the proximal **(D)** and distal **(E)** neuronal axons were quantified. **(F-G)** The total cellular peroxisome content in proximal **(F)** and distal **(G)** neurons of *acbd4/5* depleted flies normalized to that of wild-type flies. **(H-I)** The total cellular peroxisome content in proximal **(H)** and distal **(I)** neurons of ageing flies. Data were analysed by one-way ANOVA with Dunnett’s or Sidak’s multiple comparison test. ns, not significant; *, P < 0.05; ***, P < 0.001; ****, P < 0.0001. Data in graphs are expressed as mean ± SEM and n ≥ 5 flies for each group. Arrows indicate axonal localized peroxisomes. Bars: 10 µm (mains), 2.5 µm (inset).

To corroborate that the increase in axonal peroxisomes in the *acbd4/5*-depleted flies may be caused by a disruption of the Acbd4/5-Vap33 interaction and hence, peroxisome-ER contacts, we analysed peroxisomes in neurons in two fly lines with targeted *Vap33* silencing (V*ap33^RNAi.a^* and *Vap33^RNAi.b^*) (**Fig. S2A, B**). *Vap33^RNAi.a^* increased the number of peroxisomes located in proximal and distal wing axons at day 1, while *Vap33^RNAi.b^* increased axonal peroxisome numbers at 7 (distal) and 14 days (proximal and distal) compared to age-matched controls (**Fig. S2C, D**). The difference in fly age on the effect of the two *Vap33* depletion lines might be the result of a difference in RNA knockdown efficiency. Silencing of *Vap33* also did not affect the total cellular peroxisome content (**Fig. S2E, F**). Overall, depletion of *acbd4/5* or *Vap33* in *D. melanogaster* increased the number of peroxisomes located in axons, suggesting that Acbd4/5-Vap33 mediated peroxisome-ER contacts are important for peroxisome positioning in neurons.

### 2.5 Depletion of Acbd4/5 reduces lifespan of D. melanogaster but does not affect locomotion

To investigate whole-organism consequences of *acbd4/5* knockdown in neurons, we performed lifespan and rapid iterative negative geotaxis (RING) assays to assess survival and locomotor function, respectively. Pan-neuronal knockdown of *acbd4/5* with the *elav-GAL4* driver caused a significant reduction in lifespan, with a decrease in median survival to 47 days compared to 58 days in control flies (**Fig. 6A**). However, although an age associated decrease in climbing performance was observed in both groups, known to be controlled by subtle changes of the mushroom bodies and associated neurons (Sun et al., 2018), *acbd4/5^RNAi^* flies did not exhibit a reduction in climbing performance compared to control flies at all ages assayed (10 – 50 DPE) (**Fig. 6B**). It should be noted that null alleles of *acbd4/5* are fertile and viable, which could reflect metabolic compensation. However, the reduction in longevity suggests that compensation may be incomplete.

**Fig. 6.**
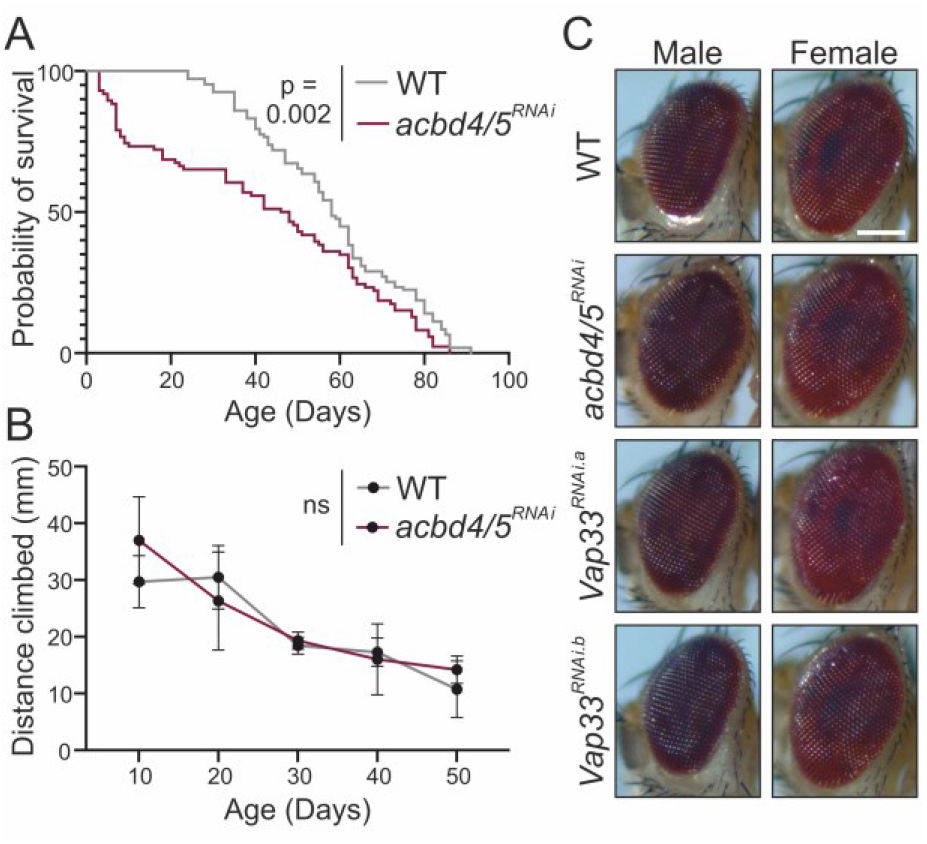
Depletion of *acbd4/5* in neurons reduces lifespan of *D. melanogaster* but does not affect locomotion. **(A)** Kaplan–Meier survival curves of control and pan-neuronal *acbd4/5^RNAi^* knockdown flies. Log-rank test, n = 85-100 flies per genotype. **(B)** Distance climbed in RING assay after 4 seconds in control and pan-neuronal *acbd4/5^RNAi^*knockdown flies at 10, 20, 30, 40 and 50 DPE. Bars represent the mean height climbed ± SD, two-way ANOVA with FDR correction, n = 3 vials of 10 flies per genotype at each age. **(C)** No overt eye phenotype was observed with *acbd4/5* or *Vap33* silencing (*GMR-GAL4*). Pictures taken at 21 DPE. Scale bar: 0.15 mm.

One of the earliest studied roles of peroxisomes in flies was in eye pigmentation. Deletion of *rosy*, which encodes for the peroxisomal enzyme xanthine dehydrogenase involved in the production of the red eye pigment, leads to a brownish eye colour (Beard and Holtzman, 1987). The Rosy eye phenotype was also observed in *Pex3* and *Pex16* mutant flies, implying that functional peroxisomes are required for development of pigment in the eye (Nakayama et al., 2011; Pridie et al., 2020). Therefore, we assessed eye phenotypes of flies with *acbd4/5^RNAi^*, *Vap33^RNAi.a^*or *Vap33^RNAi.b^* specifically driven in the eye (*GMR-GAL4*), but the *acbd4/5^RNAi^* and *Vap33^RNAi^* flies did not exhibit an overt eye phenotype compared to wild-type (**Fig. 6C**).

### 2.6 Expression of U. maydis Acbd4/5 and human ACBD4 reduces peroxisome number

Next, we explored Acbd4/5 in *U. maydis*, as a model of a highly polarised cell. As Um_Acbd4/5 lacks a FATT motif, it is interesting to compare the fungal protein to Dm_Acbd4/5 and Hs_ACBD4/ACBD5 with peroxisome-ER tethering function. To verify peroxisomal localisation of Um_Acbd4/5, we generated a mCherry-tagged variant. The localisation of the additional Acbd protein in *U. maydis*, Acbd1 (UMAG_02959), which only consists of an ACB domain without obvious targeting signals (**Fig. 1A**), was also determined. Both constructs were ectopically integrated in an *U. maydis* strain with GFP-SKL as peroxisomal marker. mCherry-Acbd1 did not colocalise with GFP-SKL and was instead observed in the cytosol of hyphal cells as anticipated (**Fig. 7A**). mCherry-Acbd4/5 revealed a clear colocalisation with GFP-SKL, confirming that Acbd4/5 is a peroxisomal protein.

**Fig. 7.**
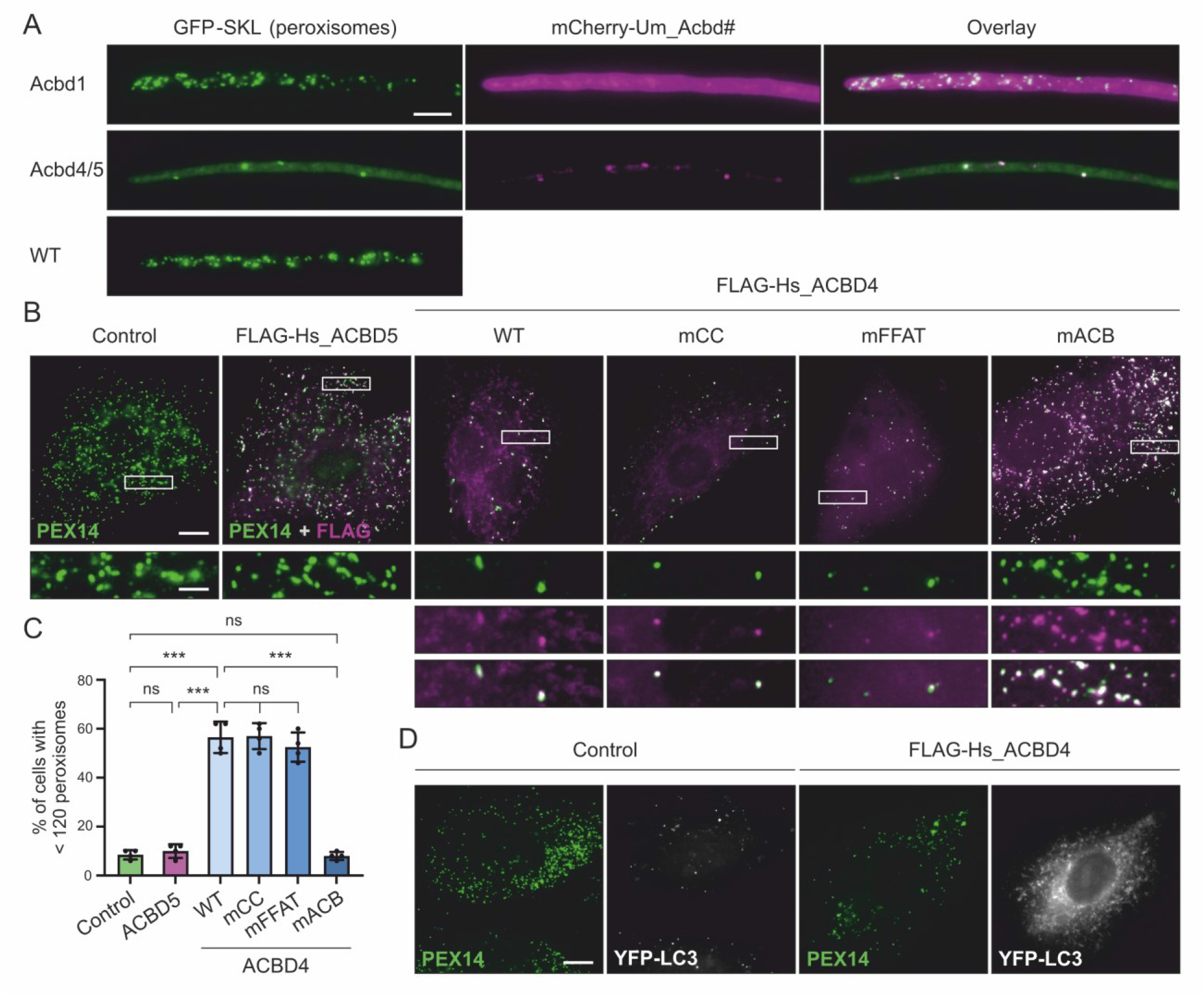
Expression of *U. maydis* Acbd4/5 and human ACBD4 reduces peroxisome number. **(A)** Subcellular localisation of *U. maydis* mCherry-Acbd1 and -Acbd4/5 in hyphal cells expressing GFP-SKL as peroxisomal marker. Bar: 5 μm. **(B)** H9C2 myoblasts transfected with human FLAG-ACBD5 or FLAG-ACBD4 WT, mCC (mutated coiled-coil domain), mFFAT or mACB were immunolabelled with PEX14 (peroxisomal marker) and FLAG antibodies. Bars: 10 µm (main), 2.5 µm (insets). **(C)** Cells with reduced peroxisome number (less than 120 peroxisomes) were quantified. Data of four replicates were analysed by one-way ANOVA with Tukey’s multiple comparisons test. n = 100 cells per condition, per replicate. ***, P < 0.001. Data are expressed as mean ± SD. **(D)** H9C2 myoblasts co-expressing FLAG-ACBD4 and autophagosomal marker YFP-LC3 were immunolabelled with PEX14.

Peculiarly, we noticed that the number of peroxisomes upon overexpression of mCherry-Acbd4/5 was reduced, accompanied by the presence of GFP-SKL staining in the cytosol (**Fig. 7A**). The ACBD4/5 homolog in the yeast *P. pastoris*, Atg37, has been implicated in regulating phagophore formation during pexophagy (Nazarko et al., 2014; Zientara-Rytter et al., 2018). Human ACBD5 has initially also been linked to pexophagy (Nazarko et al., 2014), but other studies did not reveal an involvement of ACBD5 in peroxisome degradation (Ferdinandusse et al., 2017; Yagita et al., 2017; Barone et al., 2023b). Similar observations were also recently made for *D. melanogaster* Acbd4/5 (Barone et al., 2023a). Interestingly, we noticed that overexpression of Hs_ACBD4, but not Hs_ACBD5, in H9C2 myoblasts reduced peroxisome number (**Fig. 7B, C**). While mutations in the ACBD4 coiled-coil domain or FFAT motif had no effect on the loss of peroxisomes in H9C2 cells, mutations in the ACB domain prevented the reduction in peroxisome number (**Fig. 7B, C**). Expression of ACBD4 increased fluorescence of YFP-LC3, indicating that the decrease in peroxisomes is due to autophagic processes (**Fig. 7D**). These findings indicate that *H. sapiens* ACBD4 and fungal/yeast Acbd4/5-like proteins may share a role in the regulation of peroxisome number, which may be due to an analogous development which appears to be independent of peroxisome-ER tethering.

### 2.7 Acbd4/5 deletion disrupts organelle distribution and properties in U. maydis hyphae

To examine the effect of Acbd4/5 deletion on peroxisomes in *U. maydis*, we generated an AB33 Δ*acbd4/5* hyphal strain (**Fig. S3A**). Although Um_Acbd4/5 did not bind to the ER-tethering protein Vap33 (**Fig. 4E**), GFP-SKL labelled peroxisomes surprisingly showed a redistribution, with peroxisome accumulation at the hyphal tip in Δ*acbd4/5* cells compared to an even distribution along the hyphae in wild-type cells (**Fig. 8A, B**). We previously showed that peroxisomes move along microtubules by hitchhiking on motile early endosomes in *U. maydis* (Guimaraes et al., 2015). Therefore, we decided to analyse the distribution of early endosomes in the Δ*acbd4/5* mutant. For this, we labelled early endosomes with GFP-Rab5a. Similar to the peroxisomes, the early endosomes showed accumulation at the hyphal tip in Δ*acbd4/5* cells (**Fig. 8C**). Remarkably, BODIPY 493/503 labelled lipid droplets, which also hitchhike on early endosomes (Guimaraes et al., 2015), were evenly distributed in the hyphae of both wild-type and Δ*acbd4/5* cells, while their numbers were increased in the deletion strain (**Fig. 8D, E**). We used the mitochondrial membrane potential marker TMRM to visualize mitochondria, which do not depend on early endosome transport for their motility, and observed that mitochondrial distribution was not affected upon *acbd4/5* deletion (**Fig. 8F**), implying that the absence of Acbd4/5 affects peroxisomes and early endosome positioning specifically, rather than inducing a general redistribution of organelles. However, the TMRM signal was significantly reduced, suggesting decreased mitochondrial membrane potential (**Fig. 8G**). As Acbd4/5 is an acyl-CoA binding protein, we suggest that its knockout may interrupt cellular lipid homeostasis, causing an increase in lipid droplets and potentially altering the lipid composition of membranes, which could affect organelle integrity and functional properties.

**Fig. 8.**
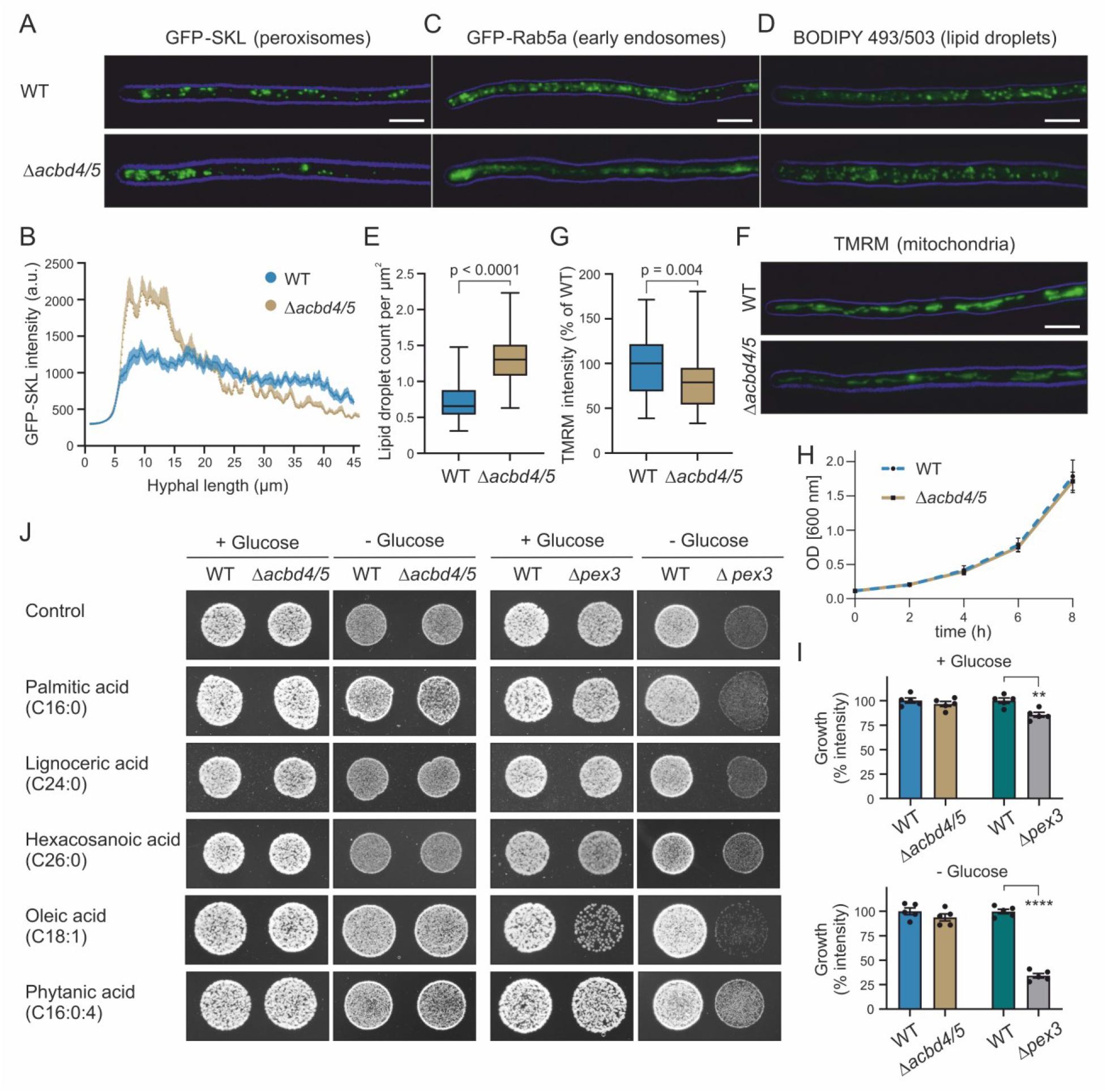
Acbd4/5 deletion disrupts organelle distribution and properties in *U. maydis* hyphae. **(A)** Peroxisome (GFP-SKL) distribution in AB33 WT and AB33 Δ*acbd4/5* hyphal cells. Bar: 5 μm. **(B)** Intensity profiles of GFP-SKL in WT and Δ*acbd4/5* cells, with the cell tips at 0 μm. Each data point represents the mean ± SEM, n = 30-32 cells of 2 experiments. **(C)** Early endosome (GFP-Rab5a) distribution in WT and Δ*acbd4/5* hyphal cells. **(D)** Lipid droplet (stained with BODIPY 493/503) distribution in WT and Δ*acbd4/5* hyphal cells. **(E)** Lipid droplet count per µm^2^ hyphae. n = 35-39 cells of 2 experiments. **(F)** Mitochondria labelled with the mitochondrial membrane potential marker TMRM (shown in green) in WT and Δ*acbd4/5* hyphal cells. **(G)** Integrated intensity measurement, normalized to mean of the WT. n = 49-56 cells of 2 experiments. Data were analysed by Mann-Whitney test. **(H)** Growth of FB1 wild-type and Δ*acbd4/5* strains in CM liquid medium complemented with 1% glucose. Data from 3 independent experiments. **(I)** Growth of wild-type (FB1 and SG200), Δ*acbd4/5* (FB1) and Δ*pex3* (SG200) strains on NM solid medium complemented with or without 1% glucose (images shown in I (control)). Data were analysed by a two-tailed unpaired t test. Data are from 5 experiments and are presented as mean ± SEM. **(J)** Strains were spotted on solid NM medium containing different fatty acids (100 μg FA/ml agar) in the presence (1000 cells) or absence (5000 cells) of 1% glucose. Images were taken after 48 h at 28°C. Quantification shown in **Fig. S3**.

### 2.8 Growth of the U. maydis Acbd4/5 deletion strain on different fatty acid substrates is unaffected

As the function of human ACBD4 and ACBD5 has been linked to peroxisomal fatty acid metabolism, we investigated the growth of an *U. maydis* FB1 Δ*acbd4/5* strain on different fatty acids (**Fig. S3B**). The growth rate of the Δ*acbd4/5* strain in liquid and on solid media was comparable to the wild-type FB1 strain (**Fig. 8H, I**). A SG200 *Δpex3* strain, which lacks peroxisomes (Camões et al., 2015), was used as a positive control. Deletion of *pex3* resulted in a reduction in growth rate in liquid (Camões et al., 2015) and on solid media compared to the SG200 wild-type strain (**Fig. 8I**). A growth defect on glucose has also been observed for *U. maydis* Δ*pex6* (peroxisome-deficient) and Δ*pex5* (restricted peroxisomal matrix protein import) strains (Freitag et al., 2012; Ast et al., 2022). In line with this, it was shown that *U. maydis* and other fungi contain several glycolytic/gluconeogenic enzymes that are dually targeted to peroxisomes and the cytosol (Idnurm et al., 2007; Freitag et al., 2012; Stiebler et al., 2014; Kremp et al., 2020; Yifrach et al., 2022).

To analyse the growth of *U. maydis* on fatty acids, wild-type, Δ*acbd4/5* and Δ*pex3* yeast-like cells were spotted on glucose-free media plates supplemented with different fatty acids (**Fig. 8J**). The integrated intensity of the individual spots was taken as measure for the ability to grow on the different carbon sources (**Fig. 8J, Fig. S3C**). We hypothesised that if *U. maydis* utilises a certain fatty acid spectrum as carbon source for growth, it would result in increased signal intensity compared to control, while if Acbd4/5/Pex3 is required, the growth of the knockout strain would be compromised compared to wild-type. To evaluate if a potential accumulation of fatty acids is toxic for fungal growth, fatty acid media supplemented with glucose was tested in parallel.

For a detailed description see **Supporting Information Fig. S3**. In summary, while a lack of peroxisomes (Δ*pex3*) impairs fungal growth on C16:0 and oleic acid, deletion of *acbd4/5* does not affect the growth of *U. maydis* on the fatty acids tested in our experimental setup.

## 3. Discussion

Our previous work revealed important functions of the human acyl-CoA binding domain proteins ACBD4 and ACBD5 in peroxisome-ER membrane contact site formation and fatty acid metabolism. Although loss of ACBD5 has been linked to human disease, our understanding of ACBD4 and ACBD5 and their specific physiological functions at the cellular and organismal level is incomplete.

### 3.1 Nomenclature and evolution

As ACBD4/5-like proteins are present in most fungi and all animals, we combined phylogenetic analyses with experimental approaches to improve understanding of their evolution and functions. Our phylogenetic analyses showed that all vertebrates possess the two paralog proteins ACBD4 and ACBD5, while invertebrates and fungi have only a single orthologous protein; following the current naming procedures, they would be called Acbp7 in *D. melanogaster* and Acbd2 in *U. maydis* (**Fig. 1A**). To avoid confusion with other ACBD proteins (e.g. human ACBD7 and ACBD2), we propose to develop a consistent nomenclature for the protein family across species. The term acyl-CoA binding domain proteins (ACBDs) would be most suitable to highlight that the proteins can perform additional functions (e.g. in contrast to Acbp, acyl-CoA binding protein). ACBD proteins can be found in all phyla and consist of four major subfamilies: (1) the small ACBD proteins, which only consist of the ACB domain (e.g. mammalian ACBD1, ACBD7, ACBD8), (2) the enoyl-CoA isomerase-coupled ACBD proteins (ACBD2), (3) the ACBDs containing protein interaction domains like ankyrin repeats or GOLD domains (ACBD3, ACBD6) and (4) the membrane-anchored ACBDs (ACBD4, ACBD5) (Islinger et al., 2020). A unifying nomenclature would therefore require extensive renaming of various historically evolved nomenclature systems found among individual plant, fungal, invertebrate and mammalian species, which is beyond the scope of this study. Hence, here we will only provisionally attempt to standardise the annotation of the tail membrane-anchored ACB domain-containing proteins, which are restricted to animals and fungi (plant ACBDs, ACBP class II-IV, have N-terminal membrane-anchoring domains and are not phylogenetically related (Lung and Chye, 2016)). Since the single membrane-anchored ACBD gene found in invertebrates and fungi does not show closer similarity to either ACBD4 or ACBD5 and, as we show here, can have distinct functions in different species, we propose to use the name Acbd4/5, as we already did in this study. In line with this, mammalian ACBD4 likely evolved from a single ancestral ACBD4/5 by a gene duplication at the base of the vertebrate branch, and consecutively reduced its ER-tethering function, while amino acid insertions in the conserved ACB domain may have changed its substrate-binding properties. Peroxisome-ER tethering is mediated through the ACBD4 and ACBD5 FFAT motif, which binds to ER-resident VAP in mammalian cells (Costello et al., 2017b; c, 2023). Interestingly, as not all ACBD4/5-like proteins contain a FFAT motif (Islinger et al., 2020), the tethering function may not be a general and ancestral function of the proteins.

### 3.2 ACBD4/5 function in D. melanogaster and U. maydis

Here, we explored the evolution and functions of ACBD4 and ACBD5 using *D. melanogaster* and *U. maydis*, as genetically-tractable model species with highly polarised cells, and showed their importance for peroxisome distribution *in vivo* in long polarized axons and hyphae, respectively. We revealed that (i) *D. melanogaster* Acbd4/5 and *U. maydis* Acbd4/5 target, like human ACBD4 and ACBD5, to peroxisomes; (ii) *D. melanogaster* Acbd4/5 binds to the ER-tethering protein Vap33 via a FFAT motif, while *U. maydis* Acbd4/5, which lacks a FFAT motif, does not; (iii) overexpression of fungal Acbd4/5 and human ACBD4 reduce peroxisome numbers while *Drosophila* ACBD4/5 and human ACBD5 induce peroxisome elongation, respectively, (iv) depletion of *acbd4/5* or *Vap33* in flies increases the number of axonal peroxisomes, and (v) *acbd4/5* deletion redistributes peroxisomes and early endosomes to the hyphal tip of fungal cells (**Fig. 9**). Our findings reveal that ACBD4/5-VAP mediated peroxisome positioning is conserved in *D. melanogaster*, and surprisingly, although the proteins do not interact in *U. maydis*, absence of Acbd4/5 does still alter peroxisome distribution.

**Fig. 9.**
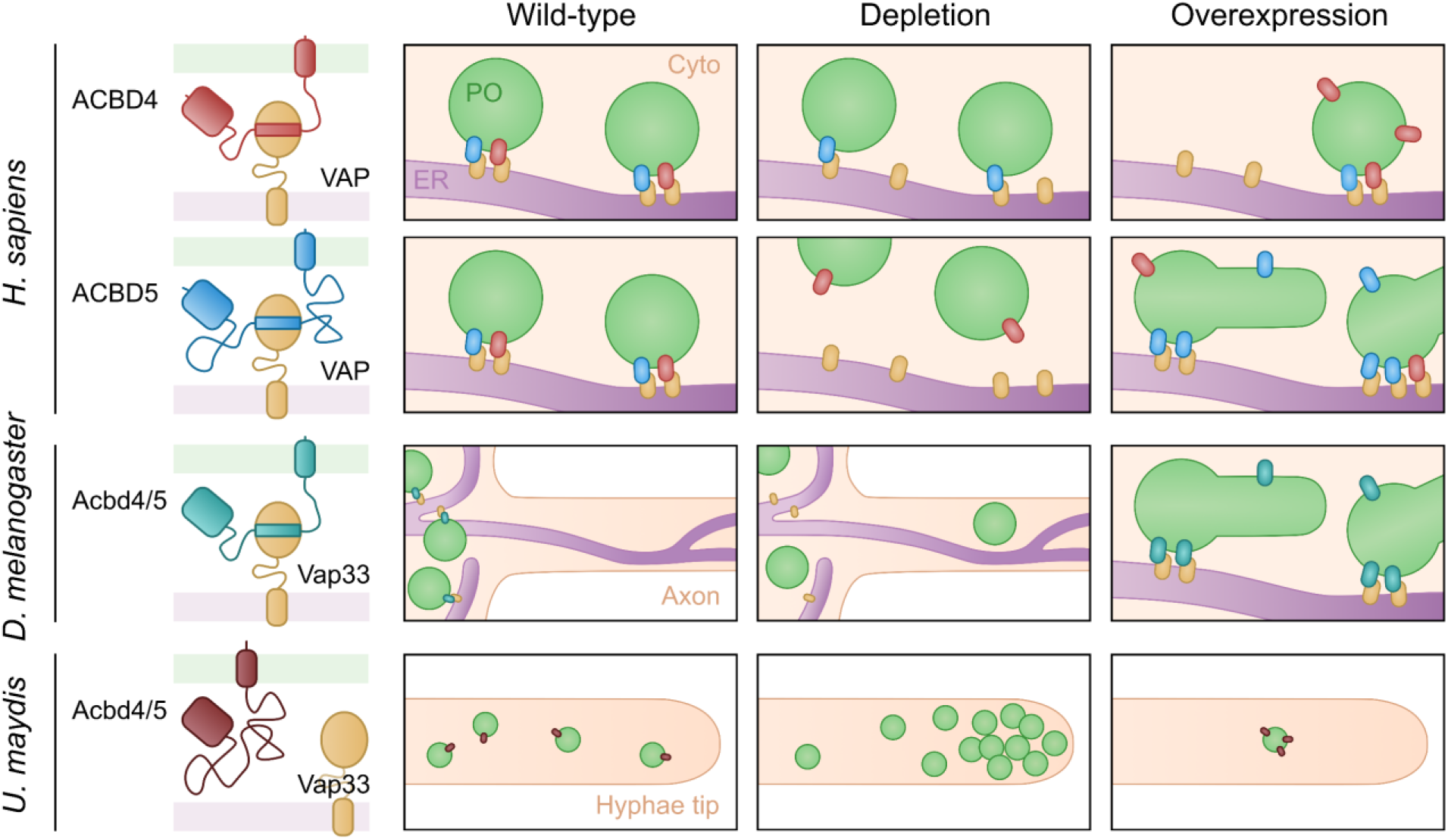
The peroxisomal ACBD4/5-like proteins in *H. sapiens*, *D. melanogaster* and *U. maydis*. Human ACBD4 and ACBD5 both bind to VAP proteins, mediating peroxisome-ER contacts. The homologs of these proteins in the fly *D. melanogaster*, Acbd4/5 and Vap33, also interact, while in the fungus *U. maydis* the homologs Acbd4/5 and Vap33 do not. In contrast to ACBD4, depletion of ACBD5 decreases peroxisome-ER contacts and hence peroxisomes become more motile. Neuronal depletion of Acbd4/5 in *D. melanogaster* enables motile peroxisomes to enter axons. Depletion of *U. maydis* Acbd4/5 also leads to a redistribution of peroxisomes, with peroxisomes accumulating at the hyphal tip. Overexpression of human ACBD4 or *U. maydis* Acbd4/5 reduces the number of peroxisomes. Overexpression of human ACBD5 induces peroxisome elongation and membrane expansion, which is required for peroxisome biogenesis. Overexpression of *D. melanogaster* Acbd4/5 seems to have a similar effect.

Organelles not only exchange metabolites, lipids and signal molecules at membrane contact sites, but also maintain their cellular positioning. For instance, contacts between the ER and mitochondria or late endosomes control their movement and distribution (Chami et al., 2008; Raiborg et al., 2015). Silencing of ACBD5 or both VAPA and VAPB results in reduced ER interaction accompanied with increased peroxisome motility in mammalian cells, implying that ER attachment restricts the movement of peroxisomes (Costello et al., 2017b; Hua et al., 2017). In line with this, depletion of *acbd4/5* or *Vap33* in *Drosophila* increased the number of peroxisomes in the axons of wing neurons without affecting overall peroxisome number. As Acbd4/5 interacts with Vap33 via its FFAT motif, we anticipate a similar tethering mechanism as in mammalian cells. Our findings suggest that disruption of the Acbd4/5-Vap33 interaction leads to peroxisome-ER detachment, with motile peroxisomes able to enter the axons.

Loss of *Vap33* has also been shown to cause relocalisation of other organelles in *Drosophila*, e.g. the Golgi apparatus which relocates from the cell body to axons and dendrites in projection neurons (Kamemura et al., 2021). Conversely, mitochondria also showed abnormal localisation after loss of *Vap33*, but they accumulated in the cell body, as opposed to being mainly distributed in dendrites and axons in wild-type neurons. Accordingly, it seems that the outcome of Vap33-mediated positioning in neurons is specific to each organelle and might be linked to the neuronal function of the organelle. Notably, as VAP proteins are involved in many ER-organelle interactions and physiological pathways, loss causes disruption of general cellular homeostasis, which could lead to indirect effects (Kors et al., 2022a).

Overexpression of ACBD5 in primary mouse hippocampal neurons has been reported to reduce peroxisome motility, with a reduction of long-range movements in neurites (Wang et al., 2018). However, overexpression of ACBD5 also increased the number of peroxisomes in neurites. Moreover, the ACBD5-induced relocalisation of peroxisomes was independent of ACBD5’s interaction with VAP, suggesting an interaction of ACBD5 with an unidentified protein that can tether, and hence redistribute peroxisomes to neurites and the plasma membrane in hippocampal neurons (Wang et al., 2018). These observations may point to cell type (and/or species) specific differences in peroxisome-ER tethering. Furthermore, cell-type specific differences in microtubule-based transport of peroxisomes may impact their redistribution after loss of ER-tethering. In mammalian cells, this depends on the adapter protein MIRO1, which can recruit kinesin/dynein motors exerting pulling forces at peroxisomes (Castro et al., 2018a; Carmichael and Schrader, 2022).

Pan-neuronal knockdown of *acbd4/5* caused a significant reduction in the survival of *D. melanogaster*, demonstrating the importance of neuronal Acbd4/5. However, we did not observe a locomotor phenotype even at 50 DPE, beyond the median lifespan of these flies. This is different from observations in patients and ACBD5 knockout mouse models, in which progressive locomotor phenotypes have been observed (Darwisch et al., 2020; Granadeiro et al., 2023). It is possible that changes in locomotion may have occurred with greater knockdown efficiency in neurons or complete ablation of the gene. ACBD5 deficient patients and mice exhibit cerebellar demyelination (Ferdinandusse et al., 2017; Yagita et al., 2017; Darwisch et al., 2020; Granadeiro et al., 2023). Whilst *D. melanogaster* do not possess myelinating oligodendrocytes, their axons are insulated by wrapping/ensheathing glial cells, ablation of which causes locomotor deficits (Kottmeier et al., 2020; Sheng et al., 2023). The contribution of glial ACBD5 to motor function could thus be a potential explanation for the lack of locomotor phenotype observed upon neuronal knockdown in *Drosophila*.

Lifespan and locomotor defects have been previously reported for *Vap33* null and ALS-associated (Vap33 P58S) mutant larvae (Chai et al., 2008; Mao et al., 2019; Karagas et al., 2022), and Vap33 P58S expressing flies (Moustaqim-Barrette et al., 2014; Sanhueza et al., 2015). Since *acbd4/5* knockdown did not affect locomotion, the effects of *Vap33* knockdown are unlikely to be caused by a reduction in Acbd4/5-Vap33 contact sites. Indeed, defects in neuromuscular junction (NMJ) boutons in the P58S mutant are attributed to a reduction in mitochondrial ATP-production, potentially via a reduction in mitochondrial Ca^2+^ uptake at mitochondria-ER contact sites (Karagas et al., 2022).

While human ACBD5 is thought to facilitate peroxisomal β-oxidation by capturing and channelling fatty acids, there is currently little information available on the metabolic function of fly Acbd4/5. A previous study that compared different *D. melanogaster* strains reported that *acbd4/5* knockdown increases levels of cuticular hydrocarbons. These lipid molecules reside in the epicuticle and have important roles in preventing desiccation and chemical signalling for social interactions (Dembeck et al., 2015). A link between peroxisomal metabolism and cuticular hydrocarbons is currently unclear.

The fungus *U. maydis* is a useful model system to uncover basic cell biological mechanisms underlying the complex regulatory networks in mammals (Steinberg and Perez-Martin, 2008). Furthermore, many peroxisomal proteins and pathways are shared between *U. maydis* and humans, including microtubule-dependent long-range transport of organelles (Camões et al., 2015). In line with this, we revealed that Um_Acbd4/5 is a peroxisomal acyl-CoA binding protein. However, in contrast to mammalian ACBD4 and ACBD5 proteins it does not contain a FFAT motif and does not interact with Vap proteins, indicating that it is not involved in peroxisome-ER tethering via Vap33 and may rather perform other (metabolic) functions. We noticed that its overexpression reduced peroxisome numbers, which was also observed in H9C2 rat myoblasts after overexpression of ACBD4. Interestingly, mutations in the ACBD4 ACB domain prevented peroxisome loss, pointing to a specific, regulatory mechanism. Notably, the ACBD4/5 homolog in the yeast *P. pastoris* (Atg37), which is required for the formation of the pexophagic receptor complex linking peroxisomes to the autophagic machinery, appears to be regulated by palmitoyl-CoA (Nazarko et al., 2014). *Drosophila* Acbd4/5 and human ACBD5 do not appear to impact peroxisome degradation (Ferdinandusse et al., 2017; Yagita et al., 2017; Barone et al., 2023a; b). We therefore suggest that *H. sapiens* ACBD4 and fungal/yeast ACBD4/5-like proteins may share a role in the regulation of peroxisome number.

Organelle positioning in the long, polarized hyphae is dependent on active diffusion and microtubule-based transport (Lin et al., 2016). As Um_Acbd4/5 does not possess a FFAT motif for Vap33-ER tethering, we hypothesised that its depletion would not impact on peroxisome distribution. Unexpectedly, however, peroxisomes were redistributed to the hyphal tips in *Δacbd4/5* cells. Notably, peroxisomes undergo kinesin and dynein-dependent transport along microtubules by hitchhiking on motile early endosomes in *U. maydis* (Guimaraes et al., 2015). Hitchhiking of peroxisomes on early endosomes was also observed in the filamentous fungus *Aspergillus nidulans* (Salogiannis et al., 2016, 2021). As early endosomes also accumulated at the tip in Δ*acbd4/5* cells, the redistribution and polar localisation of peroxisomes is likely indirect and caused by the redistribution of early endosomes. The positioning of mitochondria and lipid droplets was unaffected, although the latter also hitchhike on early endosomes. However, the increase in lipid droplet number and reduction in mitochondrial membrane potential in hyphal cells may point to a more general cellular defect and indirect effects on peroxisome/early endosome distribution, potentially caused by impaired lipid metabolism. ACBD5 deficiency results in VLCFA accumulation, and subsequent alterations in the fatty acid composition of membrane phospholipids (Darwisch et al., 2020; Granadeiro et al., 2023). These alterations could explain the observed increase in lipid droplets and changes in the mitochondrial membrane potential.

However, the yeast-like cells of the Δ*acbd4/5* strain did not show a basal growth defect in liquid and on solid media, implying normal energy production. Furthermore, the growth of the *Δacbd4/5* strain on a selection of fatty acid substrates as sole carbon source was not diminished indicating that peroxisomal fatty acid oxidation is not grossly affected. *U. maydis* possesses two fatty acid transporters (UMAG_03945, UMAG_01105) related to the mammalian peroxisomal transporters ABCD1 and ABCD2 (Camões et al., 2015). ACBD5 is supposed to capture (preferentially) VLCFA and to hand them over to ABCD1 for import into peroxisomes and subsequent β-oxidation. It is possible that alternative mechanisms for capture and hand-over of fatty acids exist in *U. maydis* or that the concentrations of fatty acids in our experimental set up are high enough to allow unassisted uptake via the *U. maydis* fatty acid transporters. It should be noted that in ACBD5 deficiency, VLCFA levels are only moderately elevated indicating that fatty acid uptake is not fully inhibited (Yagita et al., 2017; Ferdinandusse et al., 2017). Additionally, Acbd4/5 function may be specifically important in the hyphal, polarised form of *U. maydis*.

Overall, the *U. maydis* and *D. melanogaster in vivo* models provide new insights into the potential functions of ACBD4/5-like proteins as well as their evolution. They will support future studies to increase our understanding of the pathophysiological processes in ACBD5 deficient patients.

## 4. Materials and Methods

### 4.1 Bioinformatic analysis

To analyse the phylogenetic relation and structure of animal and fungal ACBD4/5-like proteins, 275 sequences from 189 species were aligned using ClustalΩ. Subsequently, phylogenetic tree construction was performed with Seaview5 (Gouy et al., 2021) using the PhyML algorithm with the following parameters: model: LG; branch support: aLRT; invariable sites: optimized; across site variation: 4 rate categories, optimized; tree searching operations: best of NNI & SPR; starting tree: BioNJ with optimized tree topology. Potential α-helical elements in the amino acid sequences were screened with PredictProtein using RePROF and ProtT5-sec (Bernhofer et al., 2021) and Jpred4 (Drozdetskiy et al. 2015), acyl-CoA binding domains with a GenomeNet motif search in PROSITE and NCBI-CDD. Coiled-coiled domains were predicted with Waggawagga using the Ncoils algorithm (Simm et al., 2015). FFAT motifs were predicted with the FFAT scoring algorithm (Murphy and Levine, 2016). Schematic diagrams of ACBD and VAP proteins in *H. sapiens*, *D. melanogaster* and *U. maydis* were created using IBS (Liu et al., 2015). For identification of protein domains, the ScanProsite tool was used (Sigrist et al., 2013). Coiled coil regions were predicted with the Marcoil (Delorenzi and Speed, 2002) and DeepCoil (Ludwiczak et al., 2019) servers using the MPI Bioinformatics Toolkit (Gabler et al., 2020). Membrane-spanning helices were detected with (Deep)TMHMM (Krogh et al., 2001; Hallgren et al., 2022).

### 4.2 Fly strains

*D. melanogaster* were housed according to rules and regulations set by the Genetically Modified Organisms committee at Cardiff University. The *D. melanogaster* strains used for experimental procedures in this study were as follows: *elav-GAL4* (Bloomington Drosophila Stock Center [BDSC]: 458); *GMR-GAL4* (1104); and *nSyb-GAL4 (51635), 10xUAS-IVS-myr::tdTomato (32221), 5xUAS-GFP::SKL (28882)*. They were crossed with the following commercially available RNAi lines and mutants: *5xUAS-CG8814^RNAi^* (67020), *CG8814^EY12940^*(transposon insertion; 21400), *5xUAS-Vap33^RNAi.a^* (27312), *5xUAS-Vap33^RNAi.b^* (77440), or control flies containing the *attP40* landing site (36304). Flies were raised under standard conditions at 25°C on cornmeal-molasses-yeast media and collected within 24 hours of eclosion. For the survival assay, female flies were housed 10 per vial in the same media and transferred to new media every 2-3 days. Surviving flies were scored at the indicated intervals.

### 4.3 Imaging of D. melanogaster wing neurons

Fly wings were imaged to examine peroxisomes in neurons according to our defined methods (Townsend et al., 2023; Maddison et al., 2023a; b; Mattedi et al., 2023). Briefly, following CO_2_ anesthesia, a wing of the fly was dissected and immediately mounted in halocarbon oil between a glass slide and a coverslip for imaging using a Zeiss Axio Examiner.Z1 microscope. A Plan-Apochromat 63x/1.40 oil DIC M27 objective (Zeiss) was used, with lasers 554 nm and 488 nm. Digital images were acquired with an Axiocam 503 mono camara using Zen 2.6 Blue (Zeiss). Z-stacks were processed and analysed as maximum projection using ImageJ. For imaging, a mixture of male and female flies was used. No sex-specific differences were observed.

To obtain a measure for the total cellular peroxisome content in neurons, a region of interest (ROI) surrounding all neurons in an image containing the tdTomato signal (used as pan-neuronal marker) was generated, and a copy of the ROI placed onto the image of the same wing area containing the GFP-SKL signal (a peroxisomal matrix marker under the same promoter as tdTomato). The integrated signal intensity of the region in both images was measured. The GFP-SKL signal was normalized to the tdTomato signal in the image.

To analyse the number of peroxisomes located in axons, peroxisomes (GFP-SKL) present in the axons (determined by tdTomato) were counted by going through the individual image planes of a Z-stack. The total number of axonal located peroxisomes in a Z-stack was normalized to the number of neuronal cell bodies (tdTomato) in the images.

### 4.4 Eye imaging

For imaging of adult eyes, flies were CO_2_ anaesthetized and frozen on dry ice. The flies were imaged using the Zeiss Stemi 508 stereo microscope with the Axiocam ERc 5s camera. Image acquisition was carried out using Zen 2.6 lite (Zeiss).

### 4.5 RING assay

The rapid iterative negative geotaxis (RING) method was used as behavioural assay for determining adult climbing ability (Gargano et al., 2005). Briefly, female flies were transferred 10 per vial to empty plastic vials without anaesthetisation. Flies were allowed to acclimatise to the vial for 10 minutes and were then dropped with equal force from a height of 100 mm. Flies were filmed as soon as the vials hit the surface and distance climbed after 4 seconds was assessed by analysing the still image at this time-point. Each vial was dropped 3 times and a mean distance climbed across the 3 repeats was calculated for each vial. Three vials of 10 flies were assessed per genotype at each age.

### 4.6 U. maydis strain generation

The *U. maydis* strains AB33_GFP-SKL, and AB33_GFP-Rab5a were published previously (Steinberg and Schuster, 2011; Schuster et al., 2011). Newly generated strains are described below. See **Table S1** for a summary of their genotypes, **Table S2** for generated and introduced plasmids and **Table S3** for details of primers used. All plasmids were generated by yeast recombination-based cloning (YRBC) in *S. cerevisiae* strain DS94 (MATα, *ura3-52, trp1-1, leu2-3, his3-111,* and *lys2-801*), which enables assembly of multiple overlapping DNA fragments in a single-cloning step (Kilaru and Steinberg, 2015). Next, plasmids were transformed into *U. maydis* strains AB33 or FB1 (Guimarães et al., 2017). Briefly, protoplasts were transformed with linearized plasmids and subsequently plated on regeneration agar plates with selectable antibiotic. Singularized transformants were screened by microscopy (for genes encoding fluorescence proteins) or Southern blotting (for generation of deletion strains).

#### AB33_mCherry-Acbd1_GFP-SKL

To visualise Acbd1 (UMAG_02959) in hyphal cells of *U. maydis*, an mCherry construct was generated. *acbd1* full length gene and terminator was amplified from *U. maydis* 521 genomic DNA (gDNA) using primers acbd1_fw and acbd1_rv (**Table S3**). Plasmid pmCherry-Acad11 (Camões et al., 2015) was digested with *Bsi*WI and *Bgl*II to remove *acad11* and replace it with *acbd1* through YRBC. The obtained pmCherry-Acbd1 plasmid was digested with *Eco*RV and integrated ectopically into the peroxisomal marker AB33_GFP-SKL strain (Steinberg and Schuster, 2011), resulting in AB33_mCherry-Acbd1_GFP-SKL.

#### AB33_mCherry-Acbd4/5_GFP-SKL

To visualise Acbd4/5 (UMAG_11226) in hyphal cells of *U. maydis*, an mCherry construct was generated. *acbd4/5* full length gene and terminator was amplified from *U. maydis* 521 genomic DNA (gDNA) using primers acbd4/5_fw and acbd4/5_rv (**Table S3**). Plasmid pmCherry-Acad11 (Camões et al., 2015) was digested with *Bsi*WI and *Bgl*II to remove *acad11* and replace it with *acbd4/5* through YRBC. The obtained pmCherry-Acbd4/5 was digested with *Eco*RV and integrated ectopically into the peroxisomal marker AB33_GFP-SKL strain (Steinberg and Schuster, 2011), resulting in AB33_mCherry-Acbd4/5_GFP-SKL.

#### AB33_ΔAcbd4/5

For deletion of *acbd4/5* in *U. maydis*, plasmid pΔAcbd4/5 was generated through YRBC. An 800-bp fragment containing part of the *acbd4/5* promoter (LF) and a 1,500-bp fragment containing downstream sequence of the *acbd4/5* gene (RF) were amplified from gDNA using primers acbd4/5_LF_fw and acbd4/5_LF_rv for LF; acbd4/5_RF_fw and acbd4/5_RF_rv for RF (**Table S3**). pΔNudE (unpublished plasmid generated with cloning vector pNEBhyg-yeast (Schuster et al., 2012)) was digested with *Bgl*II and *Bsr*GI to obtain the backbone and hygromycin resistance cassette. To generate plasmid pΔAcbd4/5, the hygromycin resistance cassette was inserted in between the promoter and terminator flank. The plasmid was digested with *Dra*I and integrated homologously into the *acbd4/5* locus of the strain AB33, resulting in AB33_ΔAcbd4/5.

#### AB33_ΔAcbd4/5_GFP-SKL

To visualise peroxisomes in AB33_ΔAcbd4/5, pGFP-SKL (Steinberg and Schuster, 2011) was linearised with DraI and integrated ectopically into the strain, resulting in AB33_ΔAcbd4/5_GFP-SKL.

#### AB33_ΔAcbd4/5_GFP-Rab5a

To visualise early endosomes in AB33_ΔAcbd4/5, pGFP-Rab5a (Schuster et al., 2011) was linearised with PciI and SspI, and integrated ectopically into the strain, resulting in AB33_ΔAcbd4/5_GFP-Rab5a.

#### FB1_ΔAcbd4/5

This strain was used for growth on different fatty acids, and deletion of *acbd4/5* in the strain FB1 was generated similarly as for AB33_ΔAcbd4/5 (see above).

### 4.7 Fungal growth conditions

For daily use, *U. maydis* strains were maintained on agar plates (1% agar (w/v), 1% glucose (w/v) in complete medium (CM; see for full composition (Holliday, 1974; Guimarães et al., 2017)). From this, liquid *U. maydis* cultures were grown overnight at 28°C in CM with 1% glucose (w/v), shaking at 200 rpm. Hyphal growth was induced by shifting to nitrate minimal medium (NM) supplemented with 1% glucose (w/v), followed by 8-14 h of growth at 28°C and 200 rpm.

### 4.8 Genomic DNA isolation and Southern blotting

To extract gDNA from *U. maydis*, cultures were grown overnight at 28°C and 200 rpm in 5 ml YEPS and centrifuged at 13,000 rpm for 1 min. The supernatant was discarded, and 0.3 g glass beads were added to the pellet, together with 400 µl of lysis buffer (2% Triton X-100, 1% SDS, 100 mM NaCl, 10 mM Tris-HCl pH 8.0, 1 mM EDTA) and 500 µl phenol-chloroform (1:1). The samples were incubated for 10 min on a Vibrax-VXR shaker (Sigma) and centrifuged at 13,000 rpm for 15 min. The upper, aqueous phase was transferred to new 1.5 ml tubes containing 1 ml of 100% ethanol and mixed by inverting. Samples were centrifuged at 13,000 rpm for 10 min, after which 500 µl 70% ethanol was added to the pellet. Samples were centrifuged at 13,000 rpm for 5 min and all ethanol was removed (residual ethanol at 55°C for 2 min). The pellet was resuspended in 50 µl ddH2O.

To confirm replacement of the *acbd4/5* gene by the hygromycin resistance cassette, analytical PCR and Southern blot analysis was performed. For PCR analysis of the gDNA, different combinations of primers binding up/downstream *acbd4/5* and in the hygromycin resistance cassette were used (Up_acbd4/5, Down_acbd4/5, Hygr, **Table S3**), so that wild-type and deletion strains could be identified based on the PCR fragment’s size or absence/presence.

For Southern blotting, gDNA was digested overnight with NcoI or PvuII. These endonucleases were chosen so that the enzyme would cut the insert and not in the wild-type *acbd4/5* locus, or in the right flank of the *acbd4/5* locus, respectively. A probe binding the left flank of the *acbd4/5* locus was generated using DIG probe labelling mix (Roche) as per the manufacturer’s instruction (primers acbd4/5_LF_fw and acbd4/5_LF_rv, **Table S3**). The digested gDNA was separated on an agarose gel. Then the gel was depurinated in 0.25 M HCl for 15 min, and subsequently neutralised in 0.4 M NaOH for 15 min. DNA was transferred from the gel to Amerhsam Hybond-NX membrane (GE Healthcare) with 0.4 M NaOH, for at least 4 h, through capillarity. The membrane was UV cross-linked to covalently bind the DNA to the membrane. The membrane was incubated with hybridization buffer (0.5 M NaPO_4_, 7% SDS) at 68°C for 30 min, and subsequently in 50 ml hybridization buffer with the probe at 68°C overnight. The following day, the membrane was washed (0.5 M NaPO_4_, 1% SDS), blocked (1% milk in DIG-buffer: 0.1 M maleic acid, 0.15 M NaCl, pH 7.5) and then incubated with Anti-Digoxigenin-AP (Roche) in 1:10 block buffer, followed by detection with Tropix CDP-Star solution (Applied Biosystems) using the G:Box Chemi (Syngene).

### 4.9 Live cell imaging of U. maydis

For microscopy of *U. maydis*, cells from a liquid culture were placed on a thin layer of 2% agarose, covered with a cover slip, and immediately observed using a motorized inverted microscope (IX83; Olympus) equipped with a PlanApo 100x/1.45 oil TIRF or UPlanSApo 60x/1.35 oil objective (Olympus) and a VS-LMS4 Laser-Merge-System with solid-state lasers (488 and 561 nm, 75 mW; Visitron System). Images were captured using a CoolSNAP HQ2 CCD camera (Photometrics). All parts of the system were controlled by the VisiView 3.3.0.4 (Visitron Systems). Z-stacks were taken using a Piezo drive (Piezosystem Jena), and processed and analysed as maximum projection using MetaMorph. ImageJ was used for image processing and overlay. Cell edges are indicated by an overlain false-colour bright-field image.

To measure peroxisome distribution (GFP-SKL) in hyphal cells, the mean fluorescent intensity over the length of individual cells was measured using the line-scan function in MetaMorph. The intensity at each data point was calculated relative to the total fluorescent intensity in the measured area.

Lipid droplets were visualized by incubating cells in the dark for 10 mins at room temperature with 5 μl/ml BODIPY 493/503 (Thermo Fisher Scientific, Loughborough, UK stock solution 1 mg/ml in DMSO). Z-stacks were taken, and the number of lipid droplets were analysed in maximum projections.

Mitochondrial membrane potential was visualized with tetramethylrhodamine methyl ester (TMRM; Thermo Fisher Scientific) as described previously (Steinberg et al., 2020). Briefly, 1 μl TMRM was added to 1 ml of cell culture and incubated in the dark at room temperature for 10 min, rotating on an SB2 Rotator. The integrated TMRM fluorescence intensity in the cells was measured and corrected by the integrated signal intensity within the same region in the image background.

### 4.10 Growth assays on fatty acids

To examine growth on different fatty acids, *U. maydis* wild-type cells (FB1, SG200), Δ*Acbd4/5* (FB1) and Δ*Pex3* (SG200) were grown overnight in CM supplemented with 1% glucose at 28°C and 200 rpm. Next day, cells were diluted to an OD_600_ of 0.15-0.20 and grown for 4 h. Cells were centrifuged for 5 min at 3,000 rpm and washed 3 times with NM. The cell suspension was diluted to 1,000,000 cells/ml and 200,000 cells/ml in NM, and 5 µl was plated on NM-agar plates with or without 1% glucose (w/v) supplemented with different fatty acids. The following fatty acids were used (from Sigma if not stated otherwise): palmitic acid (C16:0; stock solution 100 mg/ml), lignoceric acid (C24:0; 20 mg/ml), hexacosanoic acid methyl ester (C26:0; 20 mg/ml), vegetable oleic acid (18:1(n-9; Merck); 100 mg/ml) and phytanic acid (3,7,11,15-tetramethyl 16:0; 100 mg/ml). Fatty acids were dissolved in ethanol, for which palmitic acid, lignoceric acid and methyl hexacosanoic acid ester were heated to 70°C. Equal volumes of ethanol were added to the NM-agar plates as control. Plates were incubated for 48 h at 28°C and images acquired using Epson Perfection V850 Pro scanner. To quantitatively determine fungal growth, a region of interest (ROI) covering the growth area was generated and analysed using the software MetaMorph 7.8.6.0 (Molecular Devices). The integrated signal intensity of the region was measured and corrected by the integrated intensity of a same region in the image background.

### 4.11 Mammalian cell culture and transfection

COS-7 (African green monkey kidney cells, CRL-1651; ATCC) and H9C2 (rat myoblasts, embryo) (Rawlings et al., 2019) cells were cultured in Dulbecco’s Modified Eagle’s Medium (DMEM), high glucose (4.5 g/L) supplemented with 10% fetal bovine serum (FBS), 100 U/ml penicillin and 100 μg/ml streptomycin (all from Life Technologies) at 37°C with 5% CO_2_ and 95% humidity. COS-7 cells were transfected using diethylaminoethyl (DEAE)-dextran (Sigma-Aldrich) as described (Bonekamp et al., 2010). H9C2 cells were transfected by microporation (Neon Transfection system; 1400V, 20ms, 1 pulse). Cells were assayed for immunofluorescence or immunoprecipitation experiments 24-48 h after transfection.

### 4.12 cDNA constructs for studies in mammalian cells

Plasmids Myc-Hs_ACBD5 (WT; Q5T8D3-2), Myc-Hs_ACBD5 mFFAT (Y266K/C267K/S269R), FLAG-Hs_ACBD5, FLAG-Hs_ACBD4 (WT; Q8NC06-2), FLAG-Hs_ACBD4 mFFAT (F180A/D182A/E185A), FLAG-Hs_ACBD4 mCC (M244P), FLAG-Hs_ACBD4 mACB (Y43F/K47A/Y88A) and peYFP-hLC3 were published previously (Gomes and Scorrano, 2008; Costello et al., 2017c; Kors et al., 2022b; Costello et al., 2023).

N-terminally tagged constructs of *D. melanogaster* Acbd4/5 and Vap33 were generated for expression in mammalian cells. Mammalian vectors pCMV-3Tag-2a containing *acbd4/5* (CG8814), for expression of 3xMyc-Dm_Acbd4/5 (P219E or S223E), and pCMV-3Tag-1a containing *Vap33*, for expression of 3xFLAG-Dm_Vap33, were produced by GenScript (NM_134885.4 and NM_130731.3, respectively).

N-terminally tagged constructs of *U. maydis* Acbd4/5 and Vap33 were generated for expression in mammalian cells. The *acbd4/5* gene does not contain introns and therefore was amplified from plasmid pmCherry-Acbd4/5 (**Table S2**) using forward primer ATAGAATTCATGAGCAGCGCAGACGTCATCG and reverse primer TATGATATCTCAAGCGCCAGCTTCGGCGAC (5′ to 3′). The *acbd4/5* fragment was inserted into mammalian expression vector pCMV-Tag3b using EcoRI and EcoRV, resulting in Myc-Um_Acbd4/5. Mammalian vector pCMV-3Tag-1a containing *vap33* (UMAG_11696), for expression of 3xFLAG-Um_Vap33, was produced by GenScript (XM_011389787.1). All constructs produced were confirmed by sequencing (Eurofins Genomics).

### 4.13 Immunofluorescence and microscopy of cultured mammalian cells

Cells grown on glass coverslips were fixed with 4% paraformaldehyde (PFA; in PBS, pH 7.4) for 20 min, permeabilised with 0.2% Triton X-100 for 10 min, and blocked with 1% BSA for 10 min.

Blocked cells were sequentially incubated with primary and secondary antibodies (**Table S4**) for 1 h in a humid chamber at room temperature. Coverslips were washed with ddH2O to remove PBS and mounted on glass slides using Mowiol medium. Cell imaging was performed using an Olympus IX81 microscope equipped with an UPlanSApo 100x/1.40 oil objective (Olympus Optical). Digital images were taken with a CoolSNAP HQ2 CCD camera (Photometrics) and adjusted for contrast and brightness using MetaMorph 7 (Molecular Devices) or ImageJ.

### 4.14 Immunoprecipitation

Protein interactions were assayed by immunoprecipitation as previously described (Kors and Schrader, 2023). For immunoprecipitation of FLAG-Dm/Um_Vap33, the constructs mentioned in the experiments were expressed in COS-7 cells. Cells were washed in PBS and lysed in ice-cold lysis buffer (50 mM Tris-HCl pH7.4, 150 mM NaCl, 1 mM EDTA, 1% Triton X-100, mini protease inhibitor cocktail (Roche), and phosphatase inhibitor cocktail (Roche)). Insolubilized material was pelleted by centrifugation at 15,000x g. The supernatant was incubated with anti-FLAG M2 affinity gel (Sigma) at 4°C for 1 h, after which the gel was repeatedly washed with wash buffer (50 mM Tris-HCl pH7.4, 150 mM NaCl, 1 mM EDTA, 1% Triton X-100) in a rotating shaker at 4°C and by centrifugation at 5,000x g. Proteins were competitively eluted using 3X FLAG peptide (Sigma; in 10 mM Tris HCl, 150 mM NaCl, pH7.4 (TBS)).

For immunoprecipitation of Myc-Dm/Um_Acbd4/5, the constructs mentioned in the experiments were expressed in COS-7 cells. After 48 h cells were washed in PBS, and lysed in ice-cold lysis buffer (10 mM Tris-HCL pH 7.4, 150 mM NaCl, 0.5 mM EDTA, 0.5% NP-40, mini protease inhibitor cocktail (Roche), and phosphatase inhibitor cocktail (Roche)). Insolubilized material was pelleted by centrifugation at 15,000x g. The supernatant was diluted (1:2) with dilution buffer (10 mM Tris-HCL pH 7.4, 150 mM NaCl, 0.5 mM EDTA) and mixed with Myc-TRAP (ChromoTek) magnetic agarose beads and incubated for 1 h at 4°C. Beads were subsequently extensively washed with wash buffer (10 mM Tris-HCL pH 7.4, 150 mM NaCl, 0.5 mM EDTA, 0.05% NP-40) in a rotating shaker at 4°C. Proteins were eluted with Laemmli buffer for 10 min at 95°C.

Immunoprecipitates and total lysates were analysed by Western immunoblotting. Proteins were separated on 4-12% gradient precast SurePAGE gels (GenScript), and subsequently transferred to nitrocellulose membranes (Amersham Bioscience) using a semi-dry apparatus (Trans-Blot SD, Bio-Rad). Membranes were blocked in 5% dry milk (Marvel) in Tris-buffered saline with Tween-20 (TBS-T), and incubated with primary antibodies (**Table S4**), followed by incubation with horseradish peroxidase-conjugated secondary antibodies (**Table S4**) and detected with enhanced chemiluminescence reagents (Amersham Bioscience) using the G:Box Chemi (Syngene).

### 4.15 Statistical Analysis

Two-tailed unpaired t-tests were used for statistical comparisons between two groups. For experiments containing more groups, one-way ANOVA with Sidak’s post hoc test was used to determine statistical differences between the mean of selected pairs; or one-way ANOVA with Dunnett’s post hoc test was used to determine statistical differences against a control mean. Log-rank test was used for survival analysis, and two-way ANOVA with FDR correction was used for the RING assay. For these tests, data distribution was assumed to be normal but this was not formally tested. Nonparametric Mann-Whitney test was used for statistical comparison of the non-normal distributed TMRM data (at least one data set with p<0.05 in Shapiro–Wilk test). Statistical analyses were performed on GraphPad Prism (v9.4.1 and 10.2.0). Data are presented as mean ± SEM/SD. * p < 0.05, ** p < 0.01, *** p < 0.001, **** p < 0.0001.

## Acknowledgements

We would like to thank J. M. Henley and L. Lee (Bristol) for providing H9C2 cells, L. Scorrano (Padua) for providing plasmid peYFP-hLC3, and G. Steinberg (Exeter) for supporting the Ustilago-related work.

## Author Contributions (CRediT authorship contribution statement)

Suzan Kors: Conceptualization, Investigation, Formal analysis, Writing - original draft. Martin Schuster: Conceptualization, Investigation, Formal analysis, Writing - review & editing. Daniel C. Maddison: Conceptualization, Investigation, Formal analysis, Writing - review & editing. Sreedhar Kilaru: Conceptualization, Investigation, Formal analysis, Writing - review & editing. Tina A. Schrader: Investigation, Formal analysis. Joseph L. Costello: Conceptualization, Investigation, Formal analysis, Writing - review & editing. Markus Islinger: Conceptualization, Investigation, Formal analysis, Writing - original draft. Gaynor A. Smith: Conceptualization, Investigation, Formal analysis, Writing - review & editing. Michael Schrader: Conceptualization, Investigation, Formal analysis, Writing - original draft. All authors contributed to methods.

## 6. Supplementary Information

**Fig. S1.**
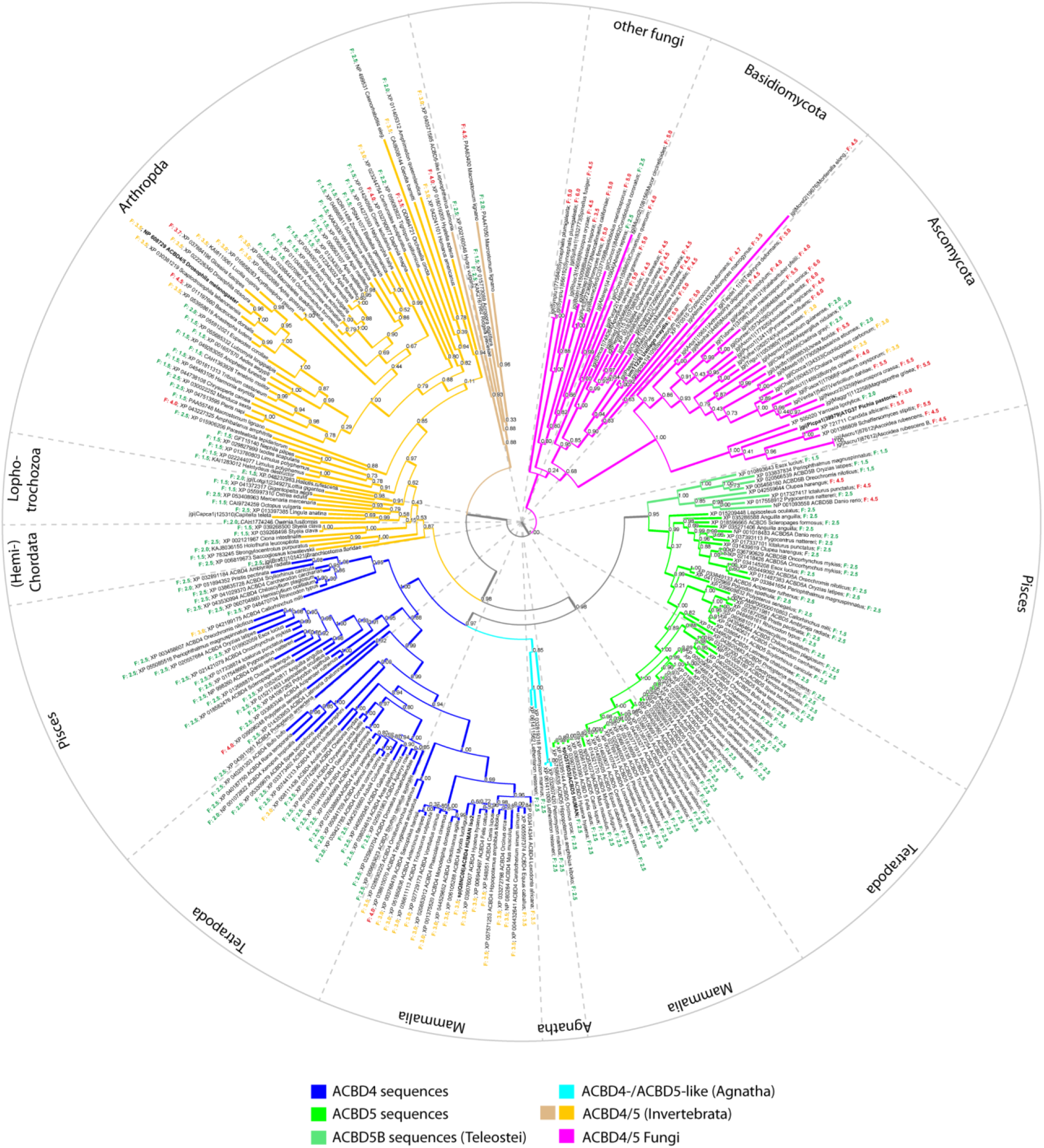
Cladogram depicting the phylogenetic relationship between ACBD4, ACBD5 and ACBD4/5 sequences from animals and fungi. Numbers at branch points show the values of branch support probabilities. F-values behind the species identifiers show values of the FFAT motif prediction algorithm (Murphy and Levine, 2016). F-values in green symbolise a potentially high affinity to the MSP domain of VAPs, and orange values indicate potentially moderate binding affinities. Red FFAT values most likely represent sequence motifs, which have affinities too low for a stable interaction with MSP domains. Note that the FFAT score does not indicate the definite binding strength. Only vertebrates possess both ACBD4 and ACBD5 genes. Note that ACBD5 sequences generally possess FFAT values indicative of stronger binding to MSP domains than ACBD4 sequences. Moreover, teleost fish exhibit a third ACBD4/5-like sequence (ACBD5B), which appears to have evolved from an additional gene duplication of the ancestral ACBD5 gene.

**Fig. S2.**
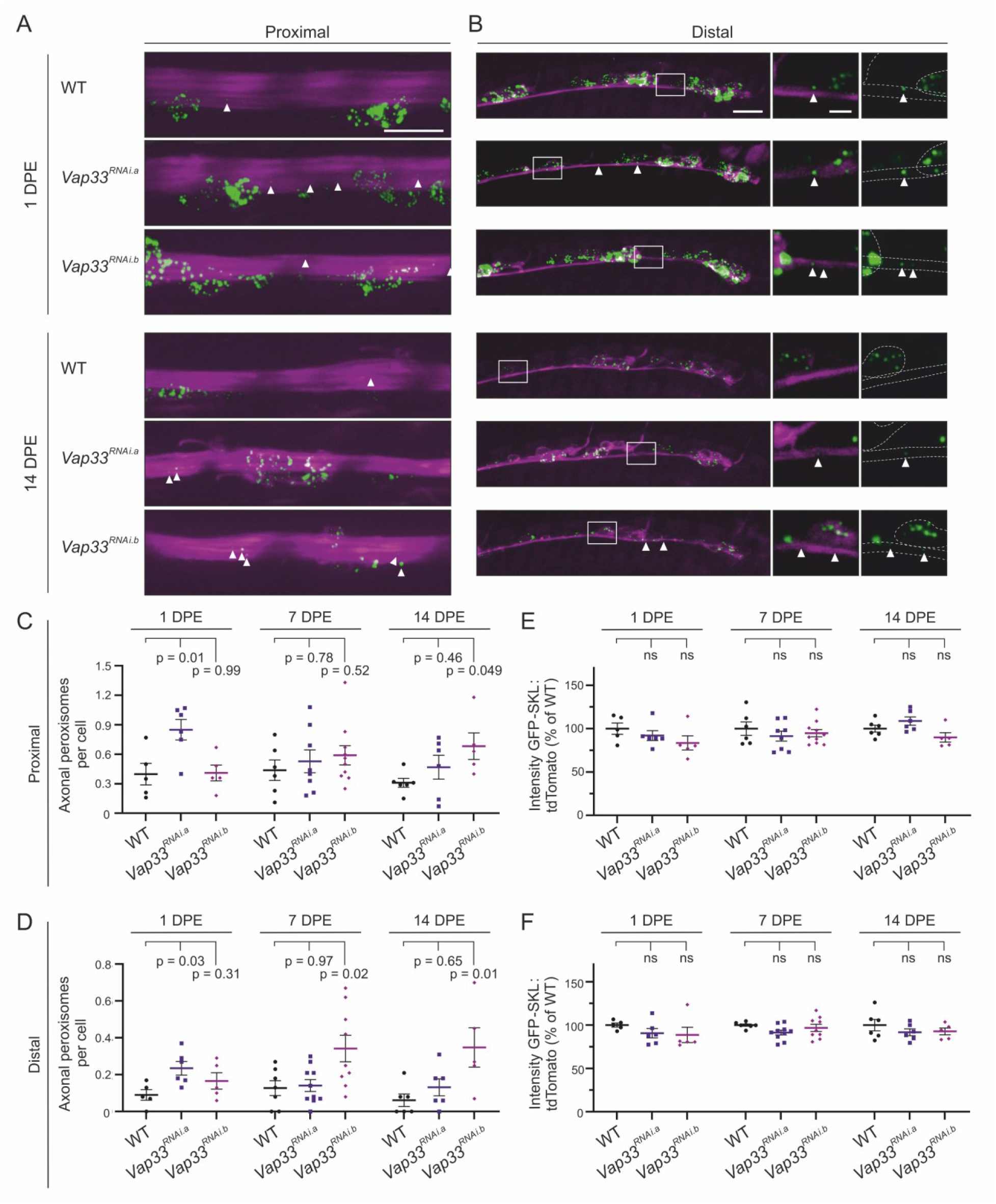
Depletion of *Vap33* in *D. melanogaster* increases the number of axonal peroxisomes. **(A-B)** Flies expressing GFP-SKL and tdTomato to label peroxisomes and the cytosol, respectively, in mature neurons were used (*nSyb-GAL4, UAS-GFP::SKL, UAS-tdTomato*). Proximal **(A)** and distal **(B)** neurons were imaged at 1 and 14 DPE in *vap33* depleted (*Vap33^RNAi.a^*and *Vap33^RNAi.b^*) flies. Insets present selected planes of the Z-projection shown in the main. **(C-D)** The number of peroxisomes located in proximal **(C)** and distal **(D)** neuronal axons at 1, 7 and 14 DPE were quantified. **(E-F)** The total cellular peroxisome content in proximal **(E)** and distal **(F)** neurons of *Vap33* depleted flies was normalized to that of wild-type flies. Data were analysed by one-way ANOVA with Dunnett’s multiple comparison test. ns, not significant. Data in graphs are expressed as mean ± SEM and n ≥ 5 flies for each group. Arrows indicate axonal localized peroxisomes. Bars: 10 µm (mains), 2.5 µm (inset).

**Fig. S3.**
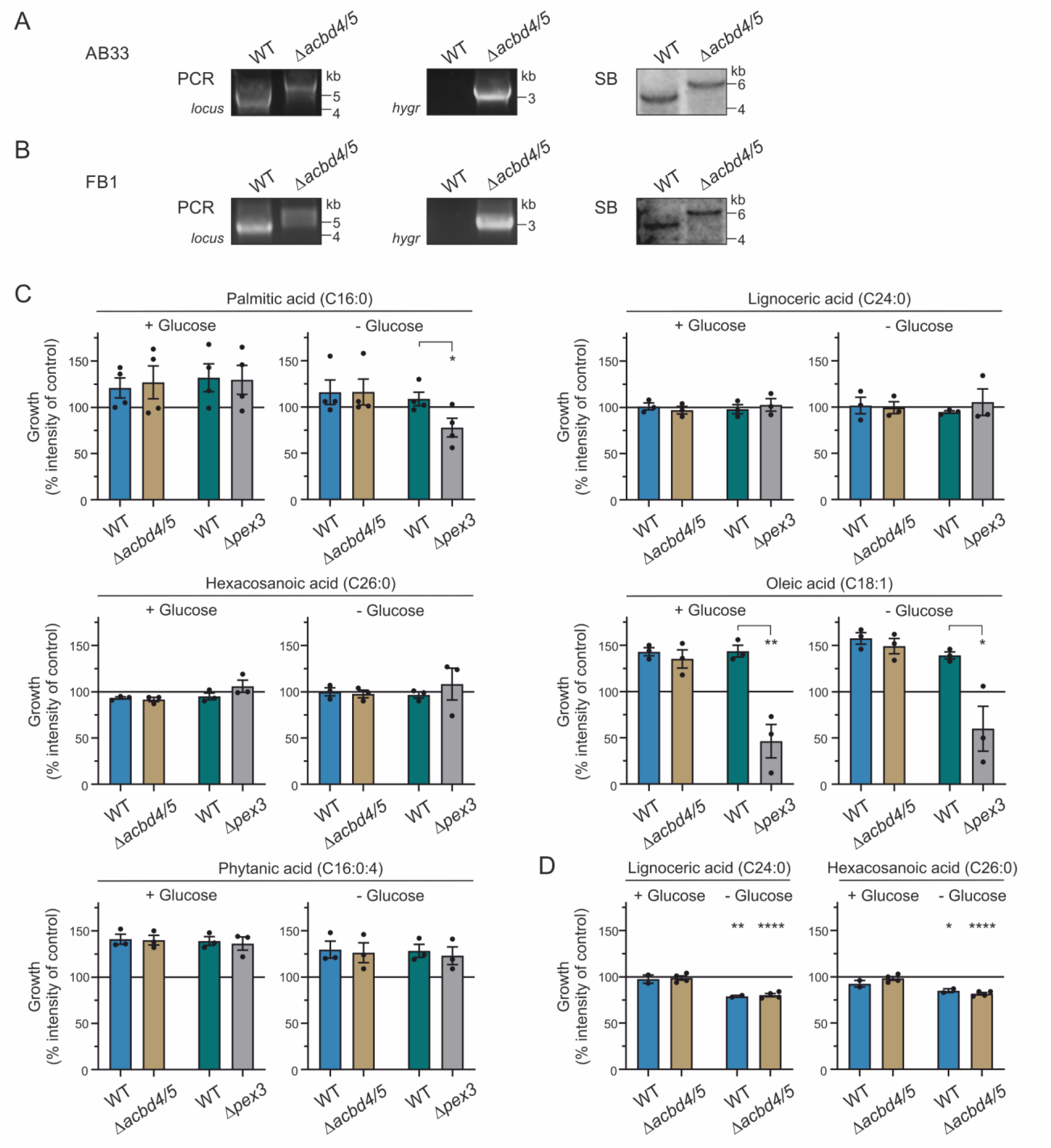
Growth of *U. maydis* Δ*acbd4/5* and Δ*pex3* strains on different fatty acids. To generate **(A)** AB33 and **(B)** FB1 *acbd4/5* deletion strains, the *acbd4/5* gene was replaced by the hygromycin resistance cassette. The deletion was confirmed by analytical PCR and Southern blot (SB). **(C)** The fungal growth of FB1 WT/Δ*acbd4/5* and SG200 WT/Δ*pex3* strains on solid NM medium containing different fatty acids (100 μg FA/ml agar), as shown in Fig. 8J, was quantified. Integrated intensity of each spot was normalized to the integrated intensity of the control (ethanol) for each experiment, which is presented by the horizontal line in each graph. **(D)** Growth of FB1 wild-type and Δacbd4/5 on solid NM medium containing 200 μg FA/ml agar of lignoceric acid (C24:0) or hexacosanoic acid (C26:0). Data were analysed by a two-tailed unpaired t test. Data are from at least three experiments and are presented as mean ± SEM. See **Supporting Information Fig. S3** for detailed information.

**Supporting Information Fig. S3. Growth of *U. maydis* Δ*acbd4/5* and Δ*pex3* strains on different fatty acids.**

Growth of the Δ*acbd4/5* mutant on palmitic acid (C16:0, a saturated LCFA) was not altered, while the growth of the Δ*pex3* mutant was significantly reduced, compared to the wild-type strains (**Fig. S3C, Fig. 8J**) (Camões et al., 2015). The growth of both Δ*acbd4/5* and Δ*pex3* on lignoceric acid (C24:0) and hexacosanoic acid (C26:0) was not compromised at the concentrations tested. As these VLCFA also did not induce increased growth compared to control – as observed with C16:0 – we doubled the concentration of C24:0 and C26:0 in the media (200 μg FA/ml agar) (**Fig. S3D**). However, this reduced the growth of both wild-type and knockout strains in the absence of glucose, suggesting that those higher concentrations lead to toxicity when there is no energy source (e.g. glucose) for detoxification. As C24:0 and C26:0 did not lead to enhanced growth, it seems that these fatty acids are not used by *U. maydis* as carbon source for growth. Interestingly, *S. cerevisiae* appears to have low peroxisomal β-oxidation rates for C24:0 and C26:0 compared to other fatty acids (e.g. <1% of the rates for C16:0 and oleic acid (C18:1)) (van Roermund et al., 2014), this might be similar in *U. maydis*.

Next, we analysed the growth of *U. maydis* on an unsaturated LCFA (oleic acid, C18:1) and a branched-chain FA (phytanic acid, C16:0:4) (**Fig. S3C**). Loss of *pex3* suppressed growth on oleic acid in both the absence and presence of glucose, suggesting that even in the presence of an alternative carbon source the growth could not be restored and hence, that the fatty acid exhibited significant toxicity (Camões et al., 2015). Impaired growth on oleic acid-containing medium was also observed with Δ*pex5*, Δ*pex6* and Δ*mfe2* mutants (Freitag et al., 2012; Kretschmer et al., 2012; Ast et al., 2022), indicating that oleic acid can only be degraded in significant amounts in peroxisomes in *U. maydis*.

However, the Δ*acbd4/5* mutant was not affected in its growth on oleic acid compared to wild-type, suggesting that Acbd4/5 does not play a critical role in its degradation (**Fig. S3C**). The Δ*acbd4/5*, Δ*pex3* and wild-type strains all showed increased growth on phytanic acid compared to control. In mammals, phytanic acid can be degraded via peroxisomal α-oxidation, or as alternative, in the ER via ω-oxidation with the resulting pristanic acid subsequently degraded in peroxisomes via β-oxidation. Although *U. maydis* does not possess a complete enzyme inventory for peroxisomal α-oxidation, it seems to utilize a similar degradation system for branched-chain fatty acids (Camões et al., 2015). As our results show enhanced growth of the peroxisome-lacking Δ*pex3* strain on phytanic acid, it suggests that peroxisomes are not essential for its degradation in *U. maydis*.

**Table S1.** *U. maydis* strains used and generated in this study.

| Strain name | Genotype | References |
| --- | --- | --- |
| AB33 | wildtype, *a2 PnarbW2 PnarbE1* | (Brachmann et al., 2001) |
| AB33_GFP-SKL | *a2 PnarbW2 PnarbE1, ble<sup>R</sup> / Potef-eGFP-SKL, cbx<sup>R</sup>* | (Steinberg and Schuster, 2011) |
| AB33_GFP-Rab5a | *a2 PnarbW2 PnarbE1, ble<sup>R</sup> / Potef-eGFP-Rab5a, nat<sup>R</sup>* | (Schuster et al., 2011) |
| AB33_mCherry-Acbd1_GFP-SKL | *a2 PnarbW2 PnarbE1, ble<sup>R</sup> / Potef-eGFP-SKL, cbx<sup>R</sup> / Potef-mCherry-Acbd1, hyg<sup>R</sup>* | This study |
| AB33_mCherry-Acbd4/5_GFP-SKL | *a2 PnarbW2 PnarbE1, ble<sup>R</sup> / Potef-eGFP-SKL, cbx<sup>R</sup> / Potef-mCherry-Acbd4/5, hyg<sup>R</sup>* | This study |
| AB33_ΔAcbd4/5 | *a2 PnarbW2 PnarbE1, ble<sup>R</sup>; ΔAcbd4/5, hyg<sup>R</sup>* | This study |
| AB33_ΔAcbd4/5_GFP-SKL | *a2 PnarbW2 PnarbE1, ble<sup>R</sup>; ΔAcbd4/5, hyg<sup>R</sup> / Potef-eGFP-SKL, cbx<sup>R</sup>* | This study |
| AB33_ΔAcbd4/5_GFP-Rab5a | *a2 PnarbW2 PnarbE1, ble<sup>R</sup>; ΔAcbd4/5, hyg<sup>R</sup> / Potef-eGFP-Rab5a, nat<sup>R</sup>* | This study |
| FB1 | wildtype, *a1b1* | (Banuett and Herskowitz, 1989) |
| FB1_ΔAcbd4/5 | *a1b1; ΔAcbd4/5, hyg<sup>R</sup>* | This study |
| SG200 | wildtype, *a1 mfa2 bW2 bE* | (Bölker et al., 1995) |
| SG200_ΔPex3 | *a1 mfa2 bW2 bE, ble<sup>R</sup>; ΔPex3, hyg<sup>R</sup>* | (Camões et al., 2015) |

*a*, *b*, mating type loci; *ble<sup>R</sup>*, phleomycin resistance; *cbx<sup>R</sup>*, carboxin resistance; *E1*, *W2*, genes of the *b* mating type locus; *eGFP*, enhanced green fluorescent protein; *hyg<sup>R</sup>*, hygromycin resistance; mCherry, monomeric cherry; *mfa2*, pheromone gene; *nar*, conditional nitrate reductase promoter; *nat<sup>R</sup>*, nourseothricin resistance; *Potef*, constitutive promoter; SKL, peroxisomal targeting signal 1; Δ, deletion; /, ectopically integrated.

**Table S2.** Plasmids used and generated in this study (*U. maydis*).

| Plasmid | Genotype | Primers | References |
| --- | --- | --- | --- |
| pGFP-SKL | *Potef-eGFP-SKL, cbx<sup>R</sup>* |  | (Steinberg and Schuster, 2011) |
| pGFP-Rab5a | *Potef-eGFP-Rab5a, nat<sup>R</sup>* |  | (Schuster et al., 2011) |
| pmCherry-Acbd1 | *Potef-mCherry-Acbd1, hyg<sup>R</sup>* | acbd1_fw<br>acbd1_rv | This study |
| pmCherry-Acbd4/5 | *Potef-mCherry-Acbd4/5, hyg<sup>R</sup>* | acbd4/5_fw<br>acbd4/5_rv | This study |
| pΔAcbd4/5 | *ΔAcbd4/5, hyg<sup>R</sup>* | acbd4/5_LF_fw<br>acbd4/5_LF_rv<br>acbd4/5_RF_fw<br>acbd4/5_RF_rv | This study |

*cbx<sup>R</sup>*, carboxin resistance; *eGFP*, enhanced green fluorescent protein; *hyg<sup>R</sup>*, hygromycin resistance; mCherry, monomeric cherry; *nat<sup>R</sup>*, nourseothricin resistance; *Potef*, constitutive promoter; SKL, peroxisomal targeting signal 1; Δ, deletion.

**Table S3.**
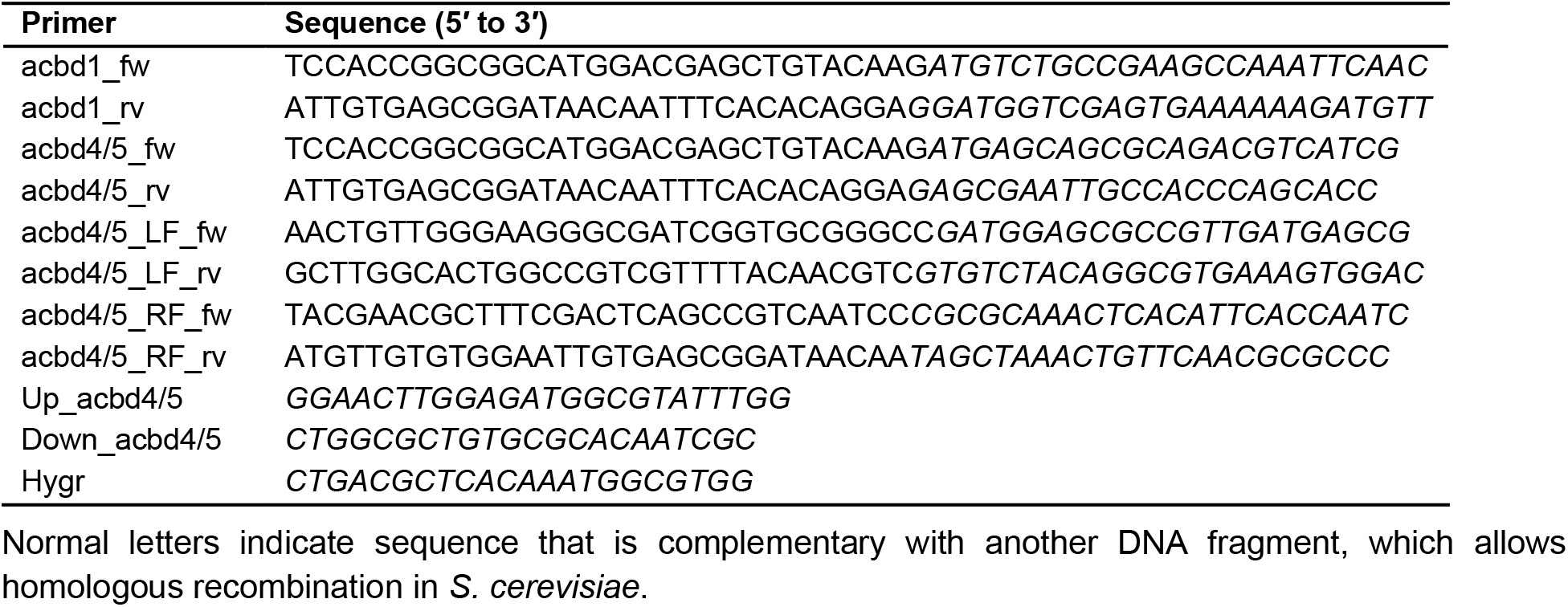
Primers used in this study (*U. maydis*).

**Table S4.** Primary and secondary antibodies used in this study.

| Antibody | Type | Dilution |  | Source |
| --- | --- | --- | --- | --- |
|  |  | WB | IMF |  |
| FLAG | mc ms | 1:2,000 |  | Sigma F3165 |
| FLAG | pc rb | 1:1,000 | 1:750 | Sigma F7425 |
| Myc | mc ms | 1:1,000 | 1:200 | Santa Cruz sc-40 |
| Myc | mc rb | 1:2,000 |  | Abcam ab9106 |
| PDI | mc ms |  | 1:100 | Thermo Fisher Scientific MA3-018 |
| PEX14 | pc rb |  | 1:14,000 | D. Crane, Griffith University, Brisbane, Australia |
| VAPB | pc rb | 1:1,000 |  | Sigma HPA013144 |
| HRP IgG | gt anti-rb | 1:10,000 |  | Bio-Rad Laboratories 170-6515 |
| HRP IgG | gt anti-ms | 1:10,000 |  | Bio-Rad Laboratories 170-6516 |
| Alexa Fluor 488 IgG | dk anti-ms |  | 1:400 | Molecular Probes A21202 |
| Alexa Fluor 488 IgG | dk anti-rb |  | 1:500 | Molecular Probes A21206 |
| Alexa Fluor 594 IgG | dk anti-ms |  | 1:1000 | Molecular Probes A21203 |
| Alexa Fluor 594 IgG | dk anti-rb |  | 1:1,000 | Molecular Probes A21207 |

Abbreviations: dk, donkey; gt, goat; HRP, horseradish peroxidase; IMF, immunofluorescence; mc, monoclonal; ms, mouse; pc, polyclonal; rb, rabbit; WB, Western blot.

**Supplemental Data 1.** Predicted ACB domains, coiled-coil structures and FFAT motifs in 275 ACBD4/5-like sequences from 189 species. The file includes ACB domain length and positioning, length and position of potential coiled-coiled motifs, and FFAT motif score, amino acid sequence and potential phosphorylation sites.

## Abbreviations

ACBD: acyl-coenzyme A binding domain containing protein
ER: endoplasmic reticulum
FFAT: two phenylalanines (FF) in an acidic tract
IP: immunoprecipitation
MSP: major sperm protein
TMD: transmembrane domain
VAP: vesicle-associated membrane protein (VAMP)-associated protein
VLCFA: very long-chain fatty acid.

## References

van Aalten, D.M.F., K.G. Milne, J.Y. Zou, G.J. Kleywegt, T. Bergfors, M.A.J. Ferguson, J. Knudsen, and T.A. Jones. 2001. Binding site differences revealed by crystal structures of Plasmodium falciparum and bovine acyl-CoA binding protein. J Mol Biol. 309:181–192. doi:10.1006/JMBI.2001.4749.

Abu-Safieh, L., M. Alrashed, S. Anazi, H. Alkuraya, A.O. Khan, M. Al-Owain, J. Al-Zahrani, L. Al-Abdi, M. Hashem, S. Al-Tarimi, M.A. Sebai, A. Shamia, M.D. Ray-Zack, M. Nassan, Z.N. Al-Hassnan, Z. Rahbeeni, S. Waheeb, A. Alkharashi, E. Abboud, S.A.F. Al-Hazzaa, and F.S. Alkuraya. 2013. Autozygome-guided exome sequencing in retinal dystrophy patients reveals pathogenetic mutations and novel candidate disease genes. Genome Res. 23:236–247. doi:10.1101/gr.144105.112.

Ast, J., N. Bäcker, E. Bittner, D. Martorana, H. Ahmad, M. Bölker, and J. Freitag. 2022. Two Pex5 Proteins With Different Cargo Specificity Are Critical for Peroxisome Function in Ustilago maydis. Front Cell Dev Biol. 10:858084. doi:10.3389/FCELL.2022.858084.

Banuett, F., and I. Herskowitz. 1989. Different a alleles of Ustilago maydis are necessary for maintenance of filamentous growth but not for meiosis. Proceedings of the National Academy of Sciences. 86:5878–5882. doi:10.1073/PNAS.86.15.5878.

Baron, M.N., C.M. Klinger, R.A. Rachubinski, and A.J. Simmonds. 2016. A Systematic Cell-Based Analysis of Localization of Predicted Drosophila Peroxisomal Proteins. Traffic. 17:536–553. doi:10.1111/TRA.12384.

Barone, F.G., M. Marcello, S. Urbé, N. Sanchez-Soriano, and M.J. Clague. 2023a. Whole organism and tissue specific analysis of pexophagy in Drosophila. bioRxiv. 2023.11.17.567516. doi:10.1101/2023.11.17.567516.

Barone, F.G., S. Urbé, and M.J. Clague. 2023b. Segregation of pathways leading to pexophagy. Life Sci Alliance. 6:e202201825. doi:10.26508/LSA.202201825.

Bartlett, M., N. Nasiri, R. Pressman, G. Bademci, and I. Forghani. 2021. First reported adult patient with retinal dystrophy and leukodystrophy caused by a novel ACBD5 variant: A case report and review of literature. Am J Med Genet. 185:1236–1241. doi:10.1002/AJMG.A.62073.

Beard, M.E., and E. Holtzman. 1987. Peroxisomes in wild-type and rosy mutant Drosophila melanogaster. Proceedings of the National Academy of Sciences. 84:7433–7437. doi:10.1073/PNAS.84.21.7433.

Bernhofer, M., C. Dallago, T. Karl, V. Satagopam, M. Heinzinger, M. Littmann, T. Olenyi, J. Qiu, K. Schütze, G. Yachdav, H. Ashkenazy, N. Ben-Tal, Y. Bromberg, T. Goldberg, L. Kajan, S. O’Donoghue, C. Sander, A. Schafferhans, A. Schlessinger, G. Vriend, M. Mirdita, P. Gawron, W. Gu, Y. Jarosz, C. Trefois, M. Steinegger, R. Schneider, and B. Rost. 2021. PredictProtein - Predicting Protein Structure and Function for 29 Years. Nucleic Acids Res. 49:W535–W540. doi:10.1093/NAR/GKAB354.

Bölker, M., S. Genin, C. Lehmler, and R. Kahmann. 1995. Genetic regulation of mating and dimorphism in Ustilago maydis. Canadian Journal of Botany. 73:320–325. doi:10.1139/B95-262.

Bonekamp, N.A., K. Vormund, R. Jacob, and M. Schrader. 2010. Dynamin-like protein 1 at the Golgi complex: A novel component of the sorting/targeting machinery en route to the plasma membrane. Exp Cell Res. 316:3454–3467. doi:10.1016/J.YEXCR.2010.07.020.

Brachmann, A., G. Weinzierl, J. Kämper, and R. Kahmann. 2001. Identification of genes in the bW/bE regulatory cascade in Ustilago maydis. Mol Microbiol. 42:1047–1063. doi:10.1046/J.1365-2958.2001.02699.X.

Camões, F., M. Islinger, S.C. Guimarães, S. Kilaru, M. Schuster, L.F. Godinho, G. Steinberg, and M. Schrader. 2015. New insights into the peroxisomal protein inventory: Acyl-CoA oxidases and -dehydrogenases are an ancient feature of peroxisomes. Biochim Biophys Acta Mol Cell Res. 1853:111–125. doi:10.1016/j.bbamcr.2014.10.005.

Carmichael, R.E., M. Islinger, and M. Schrader. 2022. Fission Impossible (?) - New Insights into Disorders of Peroxisome Dynamics. Cells. 11:1922. doi:10.3390/CELLS11121922.

Carmichael, R.E., and M. Schrader. 2022. Determinants of Peroxisome Membrane Dynamics. Front Physiol. 13:834411. doi:10.3389/FPHYS.2022.834411.

Castro, I.G., D.M. Richards, J. Metz, J.L. Costello, J.B. Passmore, T.A. Schrader, A. Gouveia, D. Ribeiro, and M. Schrader. 2018. A role for Mitochondrial Rho GTPase 1 (MIRO1) in motility and membrane dynamics of peroxisomes. Traffic. 19:229–242. doi:10.1111/TRA.12549.

Chai, A., J. Withers, Y.H. Koh, K. Parry, H. Bao, B. Zhang, V. Budnik, and G. Pennetta. 2008. hVAPB, the causative gene of a heterogeneous group of motor neuron diseases in humans, is functionally interchangeable with its Drosophila homologue DVAP-33A at the neuromuscular junction. Hum Mol Genet. 17:266–280. doi:10.1093/hmg/ddm303.

Chami, M., B. Oulès, G. Szabadkai, R. Tacine, R. Rizzuto, and P. Paterlini-Bréchot. 2008. Role of SERCA1 Truncated Isoform in the Proapoptotic Calcium Transfer from ER to Mitochondria during ER Stress. Mol Cell. 32:641–651. doi:10.1016/J.MOLCEL.2008.11.014.

Costello, J.L., I.G. Castro, F. Camões, T.A. Schrader, D. McNeall, J. Yang, E.-A. Giannopoulou, S. Gomes, V. Pogenberg, N.A. Bonekamp, D. Ribeiro, M. Wilmanns, G. Jedd, M. Islinger, and M. Schrader. 2017a. Predicting the targeting of tail-anchored proteins to subcellular compartments in mammalian cells. J Cell Sci. 130:1675–1687. doi:10.1242/jcs.200204.

Costello, J.L., I.G. Castro, C. Hacker, T.A. Schrader, J. Metz, D. Zeuschner, A.S. Azadi, L.F. Godinho, V. Costina, P. Findeisen, A. Manner, M. Islinger, and M. Schrader. 2017b. ACBD5 and VAPB mediate membrane associations between peroxisomes and the ER. Journal of Cell Biology. 216:331–342. doi:10.1083/jcb.201607055.

Costello, J.L., I.G. Castro, T.A. Schrader, M. Islinger, and M. Schrader. 2017c. Peroxisomal ACBD4 interacts with VAPB and promotes ER-peroxisome associations. Cell Cycle. 16:1039–1045. doi:10.1080/15384101.2017.1314422.

Costello, J.L., J. Koster, B.S.C. Silva, H.L. Worthy, T.A. Schrader, C. Hacker, J. Passmore, F.A. Kuypers, H.R. Waterham, and M. Schrader. 2023. Differential roles for ACBD4 and ACBD5 in peroxisome–ER interactions and lipid metabolism. Journal of Biological Chemistry. 299:105013–105014. doi:10.1016/j.jbc.2023.105013.

Darwisch, W., M. von Spangenberg, J. Lehmann, Ö. Singin, G. Deubert, S. Kühl, J. Roos, H. Horstmann, C. Körber, S. Hoppe, H. Zheng, T. Kuner, M.L. Pras-Raves, A.H.C. van Kampen, H.R. Waterham, K. V. Schwarz, J.G. Okun, C. Schultz, F.M. Vaz, and M. Islinger. 2020. Cerebellar and hepatic alterations in ACBD5-deficient mice are associated with unexpected, distinct alterations in cellular lipid homeostasis. Commun Biol. 3:1–19. doi:10.1038/s42003-020-01442-x.

Delorenzi, M., and T. Speed. 2002. An HMM model for coiled-coil domains and a comparison with PSSM-based predictions. Bioinformatics. 18:617–625. doi:10.1093/bioinformatics/18.4.617.

Dembeck, L.M., K. Bö Rö Czky, W. Huang, C. Schal, R.R.H. Anholt, and T.F.C. Mackay. 2015. Genetic architecture of natural variation in cuticular hydrocarbon composition in Drosophila melanogaster. Elife. 4:e09861. doi:10.7554/eLife.09861.001.

Færgeman, N.J., and J. Knudsen. 1997. Role of long-chain fatty acyl-CoA esters in the regulation of metabolism and in cell signalling. Biochemical Journal. 323:1–12. doi:10.1042/bj3230001.

Fan, J., X. Li, L. Issop, M. Culty, and V. Papadopoulos. 2016. ACBD2/ECI2-mediated peroxisome-mitochondria interactions in Leydig cell steroid biosynthesis. Molecular Endocrinology. 30:763–782. doi:10.1210/me.2016-1008.

Faust, J.E., A. Verma, C. Peng, and J.A. Mcnew. 2012. An Inventory of Peroxisomal Proteins and Pathways in Drosophila melanogaster. Traffic. 13:1378–1392. doi:10.1111/J.1600-0854.2012.01393.X.

Ferdinandusse, S., K.D. Falkenberg, J. Koster, P.A. Mooyer, R. Jones, C.W.T. van Roermund, A. Pizzino, M. Schrader, R.J.A. Wanders, A. Vanderver, and H.R. Waterham. 2017. ACBD5 deficiency causes a defect in peroxisomal very long-chain fatty acid metabolism. J Med Genet. 54:330–337. doi:10.1136/jmedgenet-2016-104132.

Freitag, J., J. Ast, and M. Bölker. 2012. Cryptic peroxisomal targeting via alternative splicing and stop codon read-through in fungi. Nature. 485:522–525. doi:10.1038/nature11051.

Gabler, F., S.Z. Nam, S. Till, M. Mirdita, M. Steinegger, J. Söding, A.N. Lupas, and V. Alva. 2020. Protein Sequence Analysis Using the MPI Bioinformatics Toolkit. Curr Protoc Bioinformatics. 72:e108. doi:10.1002/CPBI.108.

Gargano, J.W., I. Martin, P. Bhandari, and M.S. Grotewiel. 2005. Rapid iterative negative geotaxis (RING): A new method for assessing age-related locomotor decline in Drosophila. Exp Gerontol. 40:386–395. doi:10.1016/j.exger.2005.02.005.

Gomes, L.C., and L. Scorrano. 2008. High levels of Fis1, a pro-fission mitochondrial protein, trigger autophagy. Biochim Biophys Acta Bioenerg. 1777:860–866. doi:10.1016/J.BBABIO.2008.05.442.

Gorukmez, O., C. Havall, O. Gorukmez, and S. Dorum. 2022. Newly defined peroxisomal disease with novel ACBD5 mutation. Journal of Pediatric Endocrinology and Metabolism. 35:11–18. doi:10.1515/JPEM-2020-0352.

Gouy, M., E. Tannier, N. Comte, and D.P. Parsons. 2021. Seaview Version 5: A Multiplatform Software for Multiple Sequence Alignment, Molecular Phylogenetic Analyses, and Tree Reconciliation. Methods in Molecular Biology. 2231:241–260. doi:10.1007/978-1-0716-1036-7_15.

Granadeiro, L., V.E. Zarralanga, R. Rosa, F. Franquinho, S. Lamas, and P. Brites. 2023. Ataxia with giant axonopathy in Acbd5-deficient mice halted by adeno-associated virus gene therapy. Brain. awad407. doi:10.1093/BRAIN/AWAD407.

Guimarães, S.C., S. Kilaru, M. Schrader, and M. Schuster. 2017. Labeling of peroxisomes for live cell imaging in the filamentous fungus Ustilago maydis. Methods in Molecular Biology. 1595:131–150. doi:10.1007/978-1-4939-6937-1_13.

Guimaraes, S.C., M. Schuster, E. Bielska, G. Dagdas, S. Kilaru, B.R.A. Meadows, M. Schrader, and G. Steinberg. 2015. Peroxisomes, lipid droplets, and endoplasmic reticulum “hitchhike” on motile early endosomes. Journal of Cell Biology. 211:945–954. doi:10.1083/jcb.201505086.

Guruharsha, K.G., J.F. Rual, B. Zhai, J. Mintseris, P. Vaidya, N. Vaidya, C. Beekman, C. Wong, D.Y. Rhee, O. Cenaj, E. McKillip, S. Shah, M. Stapleton, K.H. Wan, C. Yu, B. Parsa, J.W. Carlson, X. Chen, B. Kapadia, K. Vijayraghavan, S.P. Gygi, S.E. Celniker, R.A. Obar, and S. Artavanis-Tsakonas. 2011. A Protein Complex Network of Drosophila melanogaster. Cell. 147:690–703. doi:10.1016/J.CELL.2011.08.047.

Hallgren, J., K.D. Tsirigos, M. Damgaard Pedersen, J. Juan, A. Armenteros, P. Marcatili, H. Nielsen, A. Krogh, and O. Winther. 2022. DeepTMHMM predicts alpha and beta transmembrane proteins using deep neural networks. bioRxiv. 2022.04.08.487609. doi:10.1101/2022.04.08.487609.

Hasturk, B.A., Ç. Cinar, T. Zubarioglu, S. Tiryaki-Demir, M.S. Cansever, E. Kiykim, A.K. Yigin, C. Yalcinkaya, and C. Aktuglu-Zeybek. 2024. A Novel Homozygous ACBD5 Variant in an Emerging Peroxisomal Disorder Presenting with Retinal Dystrophy and a Review of the Literature. Mol Syndromol. 15:232–239. doi:10.1159/000535534.

Helman, G., B.R. Lajoie, J. Crawford, A. Takanohashi, M. Walkiewicz, E. Dolzhenko, A.M. Gross, V.G. Gainullin, S.J. Bent, E.M. Jenkinson, S. Ferdinandusse, H.R. Waterham, I. Dorboz, E. Bertini, N. Miyake, N.I. Wolf, T.E.M. Abbink, S.M. Kirwin, C.M. Tan, G.M. Hobson, L. Guo, S. Ikegawa, A. Pizzino, J.L. Schmidt, G. Bernard, R. Schiffmann, M.S. van der Knaap, C. Simons, R.J. Taft, and A. Vanderver. 2020. Genome sequencing in persistently unsolved white matter disorders. Ann Clin Transl Neurol. 7:144–152. doi:10.1002/ACN3.50957.

Herzog, K., M.L. Pras-Raves, S. Ferdinandusse, M.A.T. Vervaart, A.C.M. Luyf, A.H.C. van Kampen, R.J.A. Wanders, H.R. Waterham, and F.M. Vaz. 2018. Functional characterisation of peroxisomal β-oxidation disorders in fibroblasts using lipidomics. J Inherit Metab Dis. 41:479–487. doi:10.1007/s10545-017-0076-9.

Holliday, R. 1974. Ustilago maydis. *In* Bacteria, Bacteriophages, and Fungi. Springer, Boston, MA. 575–595.

Hua, R., D. Cheng, É. Coyaud, S. Freeman, E. Di Pietro, Y. Wang, A. Vissa, C.M. Yip, G.D. Fairn, N. Braverman, J.H. Brumell, W.S. Trimble, B. Raught, and P.K. Kim. 2017. VAPs and ACBD5 tether peroxisomes to the ER for peroxisome maintenance and lipid homeostasis. Journal of Cell Biology. 216:367–377. doi:10.1083/jcb.201608128.

Idnurm, A., S.S. Giles, J.R. Perfect, and J. Heitman. 2007. Peroxisome function regulates growth on glucose in the basidiomycete fungus Cryptococcus neoformans. Eukaryot Cell. 6:60–72. doi:10.1128/EC.00214-06.

Islinger, M., J.L. Costello, S. Kors, E. Soupene, T.P. Levine, F.A. Kuypers, and M. Schrader. 2020. The diversity of ACBD proteins - From lipid binding to protein modulators and organelle tethers. Biochim Biophys Acta Mol Cell Res. 1867:118675. doi:10.1016/j.bbamcr.2020.118675.

Kamemura, K., C. an Chen, M. Okumura, M. Miura, and T. Chihara. 2021. Amyotrophic lateral sclerosis-associated Vap33 is required for maintaining neuronal dendrite morphology and organelle distribution in Drosophila. Genes to Cells. 26:230–239. doi:10.1111/GTC.12835.

Karagas, N.E., R. Gupta, E. Rastegari, K.L. Tan, H.H. Leung, H.J. Bellen, K. Venkatachalam, and C.-O. Wong. 2022. Loss of Activity-Induced Mitochondrial ATP Production Underlies the Synaptic Defects in a Drosophila Model of ALS. Journal of Neuroscience. JN-RM-2456–21. doi:10.1523/JNEUROSCI.2456-21.2022.

Kilaru, S., and G. Steinberg. 2015. Yeast recombination-based cloning as an efficient way of constructing vectors for Zymoseptoria tritici. Fungal Genetics and Biology. 79:76–83. doi:10.1016/J.FGB.2015.03.017.

Kors, S., J.L. Costello, and M. Schrader. 2022a. VAP Proteins – From Organelle Tethers to Pathogenic Host Interactors and Their Role in Neuronal Disease. Front Cell Dev Biol. 10:895856. doi:10.3389/FCELL.2022.895856.

Kors, S., C. Hacker, C. Bolton, R. Maier, L. Reimann, E.J.A. Kitchener, B. Warscheid, J.L. Costello, and M. Schrader. 2022b. Regulating peroxisome–ER contacts via the ACBD5-VAPB tether by FFAT motif phosphorylation and GSK3β. Journal of Cell Biology. 221:e202003143. doi:10.1083/JCB.202003143.

Kors, S., and M. Schrader. 2023. Assessing Peroxisomal Protein Interaction by Immunoprecipitation. In Peroxisomes. Methods in Molecular Biology. Humana Press Inc. 345–357.

Kottmeier, R., J. Bittern, A. Schoofs, F. Scheiwe, T. Matzat, M. Pankratz, and C. Klämbt. 2020. Wrapping glia regulates neuronal signaling speed and precision in the peripheral nervous system of Drosophila. Nat Commun. 11:4491. doi:10.1038/s41467-020-18291-1.

Kragelund, B.B., K. Poulsen, K.V. Andersen, T. Baldursson, J.B. Krøll, T.B. Neergård, J. Jepsen, P. Roepstorff, K. Kristiansen, F.M. Poulsen, and J. Knudsen. 1999. Conserved Residues and Their Role in the Structure, Function, and Stability of Acyl-Coenzyme A Binding Protein†. Biochemistry. 38:2386–2394. doi:10.1021/BI982427C.

Kremp, M., E. Bittner, D. Martorana, A. Klingenberger, T. Stehlik, M. Bölker, and J. Freitag. 2020. Non-AUG Translation Initiation Generates Peroxisomal Isoforms of 6-Phosphogluconate Dehydrogenase in Fungi. Front Cell Dev Biol. 8:251. doi:10.3389/FCELL.2020.00251.

Kretschmer, M., J. Klose, and J.W. Kronstad. 2012. Defects in mitochondrial and peroxisomal β-oxidation influence virulence in the maize pathogen Ustilago maydis. Eukaryot Cell. 11:1055–1066. doi:10.1128/EC.00129-12.

Krogh, A., B. Larsson, G. von Heijne, and E.L. Sonnhammer. 2001. Predicting transmembrane protein topology with a hidden Markov model: Application to complete genomes. J Mol Biol. 305:567–580. doi:10.1006/jmbi.2000.4315.

Lin, C., M. Schuster, S.C. Guimaraes, P. Ashwin, M. Schrader, J. Metz, C. Hacker, S.J. Gurr, and G. Steinberg. 2016. Active diffusion and microtubule-based transport oppose myosin forces to position organelles in cells. Nat Commun. 7:11814. doi:10.1038/ncomms11814.

Liu, W., Y. Xie, J. Ma, X. Luo, P. Nie, Z. Zuo, U. Lahrmann, Q. Zhao, Y. Zheng, Y. Zhao, Y. Xue, and J. Ren. 2015. IBS: an illustrator for the presentation and visualization of biological sequences. Bioinformatics. 31:3359–3361. doi:10.1093/BIOINFORMATICS/BTV362.

Ludwiczak, J., A. Winski, K. Szczepaniak, V. Alva, and S. Dunin-Horkawicz. 2019. DeepCoil—a fast and accurate prediction of coiled-coil domains in protein sequences. Bioinformatics. 35:2790–2795. doi:10.1093/BIOINFORMATICS/BTY1062.

Lung, S.C., and M.L. Chye. 2016. Deciphering the roles of acyl-CoA-binding proteins in plant cells. Protoplasma. 253:1177–1195. doi:10.1007/s00709-015-0882-6.

Maddison, D.C., F. Mattedi, A. Vagnoni, and G.A. Smith. 2023a. Clonal Imaging of Mitochondria in the Dissected Fly Wing. Cold Spring Harb Protoc. 2023:108051. doi:10.1101/PDB.PROT108051.

Maddison, D.C., F. Mattedi, A. Vagnoni, and G.A. Smith. 2023b. Analysis of Mitochondrial Dynamics in Adult Drosophila Axons. Cold Spring Harb Protoc. 2023:107819. doi:10.1101/PDB.TOP107819.

Mao, D., G. Lin, B. Tepe, Z. Zuo, K.L. Tan, M. Senturk, S. Zhang, B.R. Arenkiel, M. Sardiello, and H.J. Bellen. 2019. VAMP associated proteins are required for autophagic and lysosomal degradation by promoting a PtdIns4P-mediated endosomal pathway. Autophagy. 15:1214–1233. doi:10.1080/15548627.2019.1580103.

Mast, F.D., J. Li, M.K. Virk, S.C. Hughes, A.J. Simmonds, and R.A. Rachubinski. 2011. A Drosophila model for the Zellweger spectrum of peroxisome biogenesis disorders. Dis Model Mech. 4:659–672. doi:10.1242/DMM.007419.

Mattedi, F., D.C. Maddison, G.A. Smith, and A. Vagnoni. 2023. Live Imaging of Mitochondria in the Intact Fly Wing. Cold Spring Harb Protoc. 2023:108052. doi:10.1101/PDB.PROT108052.

Moustaqim-barrette, A., Y.Q. Lin, S. Pradhan, G.G. Neely, H.J. Bellen, and H. Tsuda. 2014. The amyotrophic lateral sclerosis 8 protein, VAP, is required for ER protein quality control. Hum Mol Genet. 23:1975–1989. doi:10.1093/HMG/DDT594.

Murphy, S.E., and T.P. Levine. 2016. VAP, a versatile access point for the endoplasmic reticulum: Review and analysis of FFAT-like motifs in the VAPome. Biochim Biophys Acta Mol Cell Biol Lipids. 1861:952–961. doi:10.1016/j.bbalip.2016.02.009.

Nakayama, M., H. Sato, T. Okuda, N. Fujisawa, N. Kono, H. Arai, E. Suzuki, M. Umeda, H.O. Ishikawa, and K. Matsuno. 2011. Drosophila Carrying Pex3 or Pex16 Mutations Are Models of Zellweger Syndrome That Reflect Its Symptoms Associated with the Absence of Peroxisomes. PLoS One. 6:e22984. doi:10.1371/JOURNAL.PONE.0022984.

Nazarko, T.Y., K. Ozeki, A. Till, G. Ramakrishnan, P. Lotfi, M. Yan, and S. Subramani. 2014. Peroxisomal Atg37 binds Atg30 or palmitoyl-CoA to regulate phagophore formation during pexophagy. Journal of Cell Biology. 204:541–557. doi:10.1083/jcb.201307050.

Neess, D., S. Bek, H. Engelsby, S.F. Gallego, and N.J. Færgeman. 2015. Long-chain acyl-CoA esters in metabolism and signaling: Role of acyl-CoA binding proteins. Prog Lipid Res. 59:1–25. doi:10.1016/j.plipres.2015.04.001.

Pandey, U.B., and C.D. Nichols. 2011. Human Disease Models in Drosophila melanogaster and the Role of the Fly in Therapeutic Drug Discovery. Pharmacol Rev. 63:411–436. doi:10.1124/PR.110.003293.

Pappaterra-Rodriguez, M.C., S.M. Muns, S.C.A. Rodríguez, G.A.R. Figueroa, N. Izquierdo, and A.L. Oliver. 2022. Variables in the ACBD5 Gene Leading to Distinct Phenotypes: A Case Report. Cureus. 14:e32930. doi:10.7759/CUREUS.32930.

Pridie, C., K. Ueda, and A.J. Simmonds. 2020. Rosy Beginnings: Studying Peroxisomes in Drosophila. Front Cell Dev Biol. 8:835. doi:10.3389/FCELL.2020.00835.

Raiborg, C., E.M. Wenzel, N.M. Pedersen, H. Olsvik, K.O. Schink, S.W. Schultz, M. Vietri, V. Nisi, C. Bucci, A. Brech, T. Johansen, and H. Stenmark. 2015. Repeated ER-endosome contacts promote endosome translocation and neurite outgrowth. Nature. 520:234–238. doi:10.1038/nature14359.

Rawlings, N., L. Lee, Y. Nakamura, K.A. Wilkinson, and J.M. Henley. 2019. Protective role of the deSUMOylating enzyme SENP3 in myocardial ischemia-reperfusion injury. PLoS One. 14:e0213331. doi:10.1371/JOURNAL.PONE.0213331.

van Roermund, C.W.T., L. Ijlst, T. Wagemans, R.J.A. Wanders, and H.R. Waterham. 2014. A role for the human peroxisomal half-transporter ABCD3 in the oxidation of dicarboxylic acids. BBA - Molecular and Cell Biology of Lipids. 1841:563–568. doi:10.1016/J.BBALIP.2013.12.001.

Rudaks, L.I., J. Triplett, K. Morris, S. Reddel, and L. Worgan. 2024. ACBD5-related retinal dystrophy with leukodystrophy due to novel mutations in ACBD5 and with additional features including ovarian insufficiency. Am J Med Genet. 194:346–350. doi:10.1002/AJMG.A.63433.

Salogiannis, J., J.R. Christensen, L.D. Songster, A. Aguilar-Maldonado, N. Shukla, and S.L. Reck-Peterson. 2021. PxdA interacts with the DipA phosphatase to regulate peroxisome hitchhiking on early endosomes. Mol Biol Cell. 32:492–503. doi:10.1091/MBC.E20-08-0559.

Salogiannis, J., M.J. Egan, and S.L. Reck-Peterson. 2016. Peroxisomes move by hitchhiking on early endosomes using the novel linker protein PxdA. Journal of Cell Biology. 212:289–296. doi:10.1083/JCB.201512020.

Sanhueza, M., A. Chai, C. Smith, B.A. McCray, T.I. Simpson, J.P. Taylor, and G. Pennetta. 2015. Network Analyses Reveal Novel Aspects of ALS Pathogenesis. PLoS Genet. 11:e1005107. doi:10.1371/JOURNAL.PGEN.1005107.

Schrader, M., M. Kamoshita, and M. Islinger. 2020. Organelle interplay - Peroxisome interactions in health and disease. J Inherit Metab Dis. 43:71–89. doi:10.1002/jimd.12083.

Schuster, M., S. Kilaru, P. Ashwin, C. Lin, N.J. Severs, and G. Steinberg. 2011. Controlled and stochastic retention concentrates dynein at microtubule ends to keep endosomes on track. EMBO J. 30:652–664. doi:10.1038/EMBOJ.2010.360.

Schuster, M., S. Treitschke, S. Kilaru, J. Molloy, N.J. Harmer, and G. Steinberg. 2012. Myosin-5, kinesin-1 and myosin-17 cooperate in secretion of fungal chitin synthase. EMBO J. 31:214. doi:10.1038/EMBOJ.2011.361.

Sheng, L., E.J. Shields, J. Gospocic, M. Sorida, L. Ju, C.N. Byrns, F. Carranza, S.L. Berger, N. Bonini, and R. Bonasio. 2023. Ensheathing glia promote increased lifespan and healthy brain aging. Aging Cell. 22:e13803. doi:10.1111/ACEL.13803.

Sigrist, C.J.A., E. De Castro, L. Cerutti, B.A. Cuche, N. Hulo, A. Bridge, L. Bougueleret, and I. Xenarios. 2013. New and continuing developments at PROSITE. Nucleic Acids Res. 41:D344–D347. doi:10.1093/NAR/GKS1067.

Simm, D., K. Hatje, and M. Kollmar. 2015. Waggawagga: comparative visualization of coiled-coil predictions and detection of stable single α-helices (SAH domains). Bioinformatics. 31:767–769. doi:10.1093/BIOINFORMATICS/BTU700.

Smith, G.A., T.-H. Lin, A.E. Sheehan, W. Van der Goes van Naters, L.J. Neukomm, H.K. Graves, D.M. Bis-Brewer, S. Züchner, and M.R. Freeman. 2019. Glutathione S-Transferase Regulates Mitochondrial Populations in Axons through Increased Glutathione Oxidation. Neuron. 103:52–65.e6. doi:10.1016/j.neuron.2019.04.017.

Steinberg, G., and J. Perez-Martin. 2008. Ustilago maydis, a new fungal model system for cell biology. Trends Cell Biol. 18:61–67. doi:10.1016/j.tcb.2007.11.008.

Steinberg, G., and M. Schuster. 2011. The dynamic fungal cell. Fungal Biol Rev. 25:14–37. doi:10.1016/J.FBR.2011.01.008.

Steinberg, G., M. Schuster, S.J. Gurr, T.A. Schrader, M. Schrader, M. Wood, A. Early, and S. Kilaru. 2020. A lipophilic cation protects crops against fungal pathogens by multiple modes of action. Nature Communications *2020 11:1*. 11:1–19. doi:10.1038/s41467-020-14949-y.

Stiebler, A.C., J. Freitag, K.O. Schink, T. Stehlik, B.A.M. Tillmann, J. Ast, and M. Bölker. 2014. Ribosomal Readthrough at a Short UGA Stop Codon Context Triggers Dual Localization of Metabolic Enzymes in Fungi and Animals. PLoS Genet. 10:e1004685. doi:10.1371/JOURNAL.PGEN.1004685.

Sun, J., A.Q. Xu, J. Giraud, H. Poppinga, T. Riemensperger, A. Fiala, and S. Birman. 2018. Neural control of startle-induced locomotion by the mushroom bodies and associated neurons in Drosophila. Front Syst Neurosci. 12:6. doi:10.3389/FNSYS.2018.00006.

Townsend, L.N., H. Clarke, D. Maddison, K.M. Jones, L. Amadio, A. Jefferson, U. Chughtai, D.M. Bis, S. Züchner, N.D. Allen, W. Van der Goes van Naters, O.M. Peters, and G.A. Smith. 2023. Cdk12 maintains the integrity of adult axons by suppressing actin remodeling. Cell Death Discov. 9:1–12. doi:10.1038/s41420-023-01642-4.

Wanders, R.J.A., H.R. Waterham, and S. Ferdinandusse. 2018. Peroxisomes and their central role in metabolic interaction networks in humans. In Proteomics of Peroxisomes. Subcellular Biochemistry. L.A. del Río and M. Schrader, editors. Springer, Singapore. 345–365.

Wang, Y., J. Metz, J.L. Costello, J. Passmore, M. Schrader, C. Schultz, and M. Islinger. 2018. Intracellular redistribution of neuronal peroxisomes in response to ACBD5 expression. PLoS One. 13:e0209507. doi:10.1371/journal.pone.0209507.

Yagita, Y., K. Shinohara, Y. Abe, K. Nakagawa, M. Al-Owain, F.S. Alkuraya, and Y. Fujiki. 2017. Deficiency of a retinal dystrophy protein, acyl-coa binding domain-containing 5 (ACBD5), impairs peroxisomal β-oxidation of very-long-chain fatty acids. Journal of Biological Chemistry. 292:691–705. doi:10.1074/jbc.M116.760090.

Yifrach, E., D. Holbrook-Smith, J. Bürgi, A. Othman, M. Eisenstein, C. Wt Van Roermund, W. Visser, A. Tirosh, M. Rudowitz, C. Bibi, S. Galor, U. Weill, A. Fadel, Y. Peleg, R. Erdmann, H.R. Waterham, R.J.A. Wanders, M. Wilmanns, N. Zamboni, M. Schuldiner, and E. Zalckvar. 2022. Systematic multi-level analysis of an organelle proteome reveals new peroxisomal functions. Mol Syst Biol. 18:e11186. doi:10.15252/MSB.202211186.

Zientara-Rytter, K., K. Ozeki, T.Y. Nazarko, and S. Subramani. 2018. Pex3 and Atg37 compete to regulate the interaction between the pexophagy receptor, Atg30, and the Hrr25 kinase. Autophagy. 14:368–384. doi:10.1080/15548627.2017.1413521.

