## Supplementary figures and images for "New insights into the functions of ACBD4/5-like proteins using a combined phylogenetic and experimental approach across model organisms"

### Figure S1

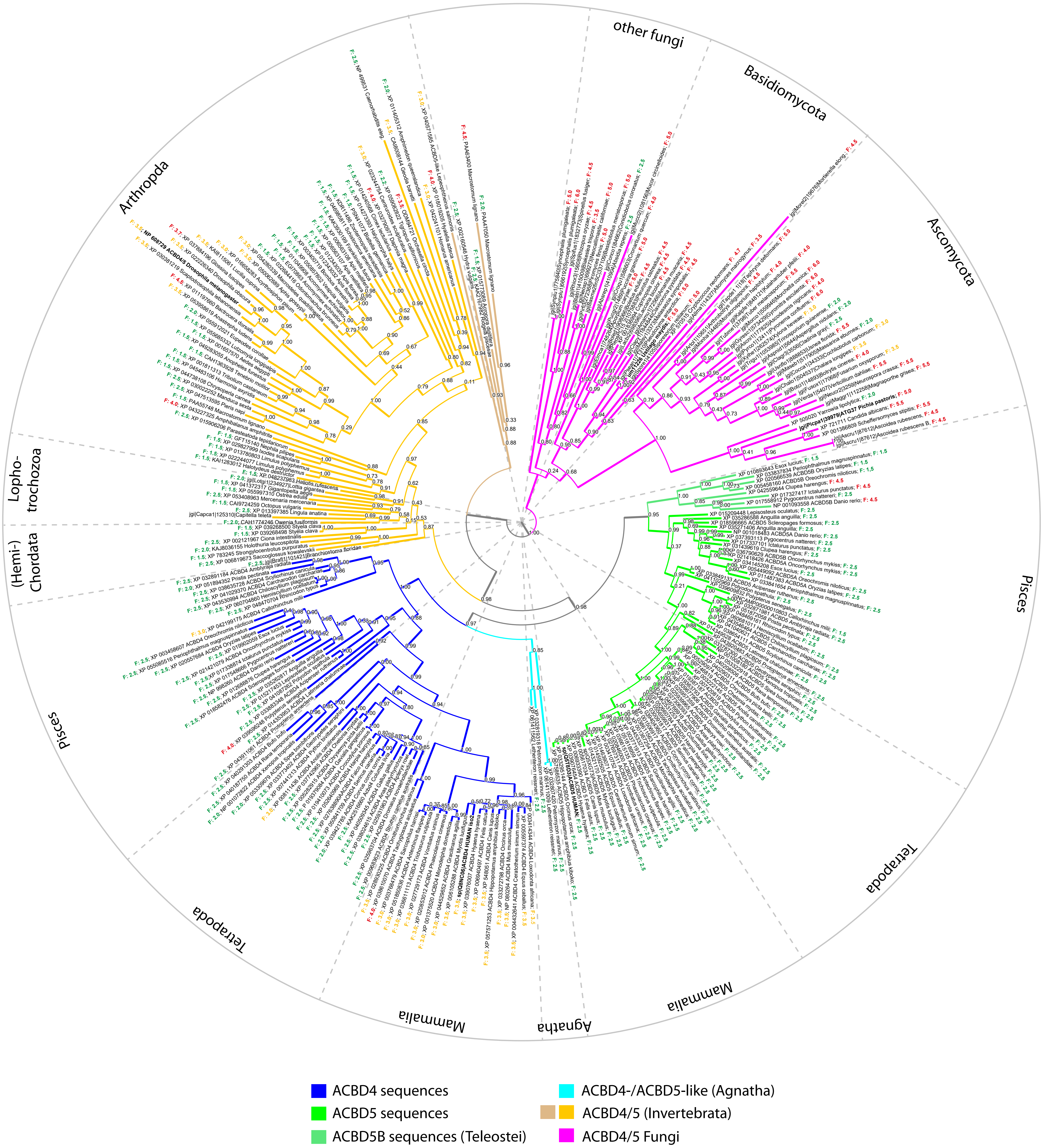
